# Targeted quantification assays for DNA repair and handling proteins and interactions in Huntington’s disease models

**DOI:** 10.64898/2026.07.28.741339

**Authors:** Todd M. Greco, Josiah E. Hutton, Joshua L. Justice, Tavis J. Reed, Thomas F. Vogt, Brinda C. Prasad, Ileana M. Cristea

## Abstract

Huntington’s disease (HD) is a life-altering genetic neurodegenerative disorder, with cognitive, motor, and psycho-social effects that have consequential impacts on the individuals and their families. While current treatments improve disease symptoms, there are no FDA-approved therapies that prevent disease progression. Converging lines of evidence from human GWAS and mouse models point to DNA repair and handling (R/H) proteins as promising therapeutic targets due to their ability to modulate somatic expansion of the CAG repeat of HTT. The roles of DNA R/H HD modulator proteins are incompletely understood, in part, due to their relatively low cellular abundance and technical challenges in quantification. Here, we developed and validated targeted mass spectrometry assays quantifying DNA R/H proteins, spanning functions in mismatch repair, Fanconi anemia, and transcriptional regulation, using complementary workflows for timsTOF and Orbitrap platforms. We built species-specific experiment spectral libraries that outperformed in silico libraries for target detection. Applying this pipeline to an HTT-Q140 knock-in mouse HD model, we observed that DNA R/H protein abundances were largely unchanged in HD mice, while HTT and HAP40 showed increased nuclear association with disease progression. To facilitate translational research applications, we further developed a stable isotope dilution assay for absolute quantification of 11 human mismatch repair-associated proteins and generated an HTT knock-out human neuroblastoma cell line. Additionally, we used thermal proximity coaggregation profiling to characterize the endogenous interactomes of MMR proteins. We observed that HTT KO caused proteome down-regulation in selected DNA R/H proteins and reshaped the MMR protein interactome, with the most pronounced changes observed for MLH1 and PMS1 interactions. Overall, we established a validated, transferable assay for quantifying DNA R/H proteins in perturbation studies using human and mouse HD model systems and provide evidence that HTT influences the abundance and interaction landscape of proteins central to CAG repeat instability.

## Introduction

Huntington’s disease (HD) is a devastating genetic neurodegenerative disorder characterized by progressive decline in motor, cognitive and emotional aspects of an affected individual. In contrast to neurodegenerative diseases such as Alzheimer’s and Parkinson’s, HD is a monogenic triplet repeat disorder and part of the family of polyglutamine repeat disorders (1). This disorder is autosomal dominant, where one copy of the huntingtin (HTT) gene contains more CAG repeats in exon 1 than the WT allele. If the individual inherits an HTT gene with a CAG repeat length above a critical threshold, typically ∼40 repeats, the penetrance is essentially complete. In individuals with HD, the expanded CAG repeat sequence is somatically unstable and prone to further expansion in vulnerable cells; thus, somatic expansion is currently viewed as causal to disease progression and contribute to the typical mid-life onset of symptomatic disease (2–7).

Currently approved treatments for HD are primarily for managing chorea symptoms; however, several promising avenues are being pursued for the development of HD therapies, with approaches that are in pre-clinical development as well as highly promising Phase I/II clinical trials (8). Several attractive drug target candidates have been generated from GWAS studies of individuals with HD, identifying DNA repair and handling (R/H) genes that modulate age of onset or CAG repeat expansion in blood (2, 3, 9, 10). Subsequently, mechanistic pre-clinical in vitro and animal studies have shown that manipulation of the expression and/or activity of mismatch repair (MMR) proteins can alter HD pathobiology and disease trajectory, underscoring their therapeutic relevance. For example, in HD knock-in mouse models, *Msh3* is required both for somatic CAG expansion and for the emergence of an early striatal disease phenotype (11). Naturally occurring *Msh3* polymorphisms that alter protein levels track with relative CAG instability (12). Consistent with this observation, in human cells, inactivation or overexpression of *MSH3* led to defective or elevated CTG•CAG repeat expansion activity, respectively (13). Subsequent work has established that siRNA-mediated *Msh3* knockdown dose-dependently slows expansion (14), while *Msh3* knockout halts somatic expansion and corrects synaptic and locomotor deficits in Q140 mice (15). Interestingly, phenotypic benefit may be contingent on repeat length, as *Msh3* ablation in zQ175 mice prevented expansion without improving striatal transcriptional dysregulation or HTT aggregation (16) Loss of *Mlh1*, *Mlh3*, or *Pms1* in HD mice similarly restricts repeat expansion (15, 17, 18), whereas in human cell and patient-derived neuronal models, FAN1 knockdown enhances CAG expansion, while its overexpression suppresses it (19, 20). Taken together, these reports unequivocally demonstrate that mismatch repair is fundamental to HD pathogenesis and that somatic CAG instability is a major driver of HTT toxicity and disease progression (7). Given the therapeutic potential of abrogating somatic expansion to slow or halt disease progression, there is high interest in characterizing the pathway level effects of manipulating MMR proteins (21). Towards this goal, the development and validation of precise and accurate multiplexed assays to measure their expression in disease-relevant tissues and cells is vital.

Integration of the accumulated knowledge of protein interaction studies with multiomics and GWA studies highlights an important role for proteins in DNA R/H and transcriptional regulatory pathways in HD natural history. Moreover, variation in abundance of DNA repair and handling factors (e.g., MutSβ) between different tissues/cell types is associated with and/or may directly contribute to rates of somatic instability (5). Specifically, recent studies in HD patient brain tissues suggests that MutSβ expression levels and transcriptional regulation may be cell type dependent, with higher levels of MutSβ in vulnerable neurons (5, 22). However, it is not known if the tissue abundance of other proteins in this pathway, including the GWAS targets, MutL proteins and FAN1, would also support this hypothesis. There has been no systematic study to quantify these proteins with high reproducibility and precision. Broadly, by developing and deploying MS-based assays that measure the dynamic regulation of proteins in DNA R/H pathways, the relationship between somatic instability and DNA R/H protein abundances in humans and mouse models can be illuminated.

Quantification of mRNA levels for MMR targets can often be determined for most biological samples (23). However, measuring their steady-state proteins levels is more challenging, often due to issues with antibody specificity and/or detection being at the lower range of quantification (24). Also, assays require sufficient sensitivity and accuracy to quantify changes in protein levels with HD relevant perturbations, such as expression of mutant huntingtin in HD mouse models (25) or reductions of huntingtin in cells (26). Therefore, development of targeted mass spectrometry assays that reproducibly measure high-responding signature peptides derived from the target proteins is desirable. These assays can be deployed on multiple instrument platforms for a broad range of sample types. They can be used for relative quantification of the same proteins across multiple samples (27), most commonly using label-free quantification (28, 29). Yet, label-free approaches can only provide estimates of absolute amounts/concentrations (24, 30). Moreover, their accuracy may be reduced when the proteomes being compared have different compositions and dynamic ranges, such as between different cell types or organs. In these cases, using stable isotope dilution mass spectrometry assays is the gold-standard for obtaining absolute quantification measurements that are broadly comparable across diverse cell and tissue types (31, 32). Independent of assay type, they can be transferred to other laboratories or core facilities with routine adjustments based on instrumentation. Historically, targeted assay development has focused on triple-quadrupole and Q-Exactive instrument platforms using selected or parallel reaction monitoring (SRM/PRM). However, for the latest generation of instruments with ultra-fast scanning rates, such as timsTOF and Astral mass analyzers, a comparison of quantitative performance between non-targeted (DIA) and targeted (PRM) acquisition has not been fully explored. Overall, development of quantitative proteins assays that are inherently multiplexed and tailored to the latest generation of instrumentation will enable reproducible measurement of DNA repair and handling protein abundances across different rodent and mammalian HD models.

## Experimental Procedures

### Experimental Design and Statistical Rationale

The total number of samples for spectral library generation and quantitative DIA and PRM experiments have been itemized and described in Supplemental Data 1. Samples prepared for statistical analysis were analyzed in at least three biological replicates per condition using GraphPad Prism 11. For processing of raw DDA and DIA data, FragPipe (ver. 22 or 24) (33, 34) was used for spectral processing, scoring identification, and FDR calculation, while DIA-NN (ver. 1.9 or 2.3.2) (35) or Skyline (ver. 26)(36) was used to perform Tier 3 MS2-based quantification (see sections below for experiment-specific processing information). For statistical analyses of proteome-level DIA data, DIA-NN protein group reports were pre-processed and analyzed for differential abundance using DEqMS (37). Differential detections were assigned by magnitude of change and Benjamini-Hochberg corrected p-value (FDR) thresholds. For statistical analysis of targeted DIA and PRM quantification experiments (< 40 proteins), peptide quantitative values were transformed, row normalized, averaged within proteins, and unpaired t-tests were performed (Gaussian data distribution and uniform standard deviation was assumed within proteins across conditions). For absolute quantification assays (Tier 2), light and heavy synthetic peptides were used for RT alignment and construction of external standard curves. Curves were constructed using linear regression and 1/Y weighting of light/heavy peptide ratios (calculated from fragment XICs) versus known on-column peptide quantity (n = 3 replicates per quantity) and were used to interpolate unknown peptide concentrations (n = 3 for standards and unknowns). Protein concentrations were calculated as the average of all peptide values and replicates (n = 3, +/- SEM). For spectral library generation, samples were prepared in three biological replicates, except for samples that were fractionated offline, which were prepared in a single replicate as they were used for qualitative purposes.

### Cell culture

BE(2)-C cells were purchased from ATCC (#CRL-2268) and cultured according to the product documentation in a 1:1 mixture of Eagle’s Minimal Essential Media and F12 Medium. For knock-out experiments, CRISPR was performed as described with the TrueCut system and CRISPRmax transfection reagent (38) gRNA was transfected into non-differentiated BE(2)-C cells in a 1:1 ratio with Cas9.

### Sample preparation for label-free DIA-PASEF in BE(2)-C cells

Cells were grown to 70% confluence, trypsinized, and washed three times with ice cold 1xPBS. Samples were lysed with 5% SDS supplemented with 1x HALT protease inhibitor cocktail (Thermo Scientific). Samples were then sonicated in a cup horn sonicator for 15 pulses and heated for 5 minutes at 95°C. Samples were then pelleted by centrifugation at 2,000 x g for 5 minutes at room temperature. Samples were then reduced and alkylated with 25 mM TCEP and 50 mM chloroacetamide for 20 minutes at 70°C. Samples were subjected to methanol/chloroform precipitation, and the resulting protein pellet was resuspended in 100 mM HEPES, pH 8.3, where the protein concentration was estimated with the BCA protein assay (Thermo Scientific). An aliquot of 50 μg of protein was removed and subjected to overnight trypsin with MS grade trypsin (Thermo Scientific) digestion at a 1:50 trypsin:protein ratio at a protein concentration of 0.5 mg/mL at 37°C with shaking at 600 rpm. The resulting peptides were desalted off-line by stage tip desalting with SDB-RPS Extraction Disks (Fisher Scientific), and peptide concentration was estimated by nanodrop measurement using the Scopes method. Peptides were resuspended in 0.015% n-dodecyl β-D-maltoside (DDM), 4% UHPLC-MS quality acetonitrile (Thermo Scientific), and 0.1% LC-MS quality formic acid (FA) in UHPLC-MS Water (Thermo Scientific) at a concentration of 100 ng/µL.

### DIA-PASEF MS/MS acquisition for relative quantification in BE(2)-C cells

LC-MS/MS analysis was performed with a nanoElute 2 attached to a timsTOF Ultra (Bruker) system. The mobile phases were 0.1% LC-MS FA in 99.9% UHPLC-MS water (buffer A) and 0.1% FA in 99.9% UHPLC-MS acetonitrile (Thermo, buffer B). A one column method with a PepSep Ultra (25 cm x 75 μm x 1.5 µm) C18 HPLC column (Bruker) and a 10 μm emitter (Bruker) attached to a CaptiveSpray Ultra source was used for LC-MS analysis. A linear 20 minute gradient of 3% to 34% buffer B at a flow rate of 200 nL/min was used for peptide separation. For data independent acquisition parallel accumulation serial fragmentation (DIA-PASEF), the MS1 settings were set to start at 100 *m/z* and end at 1,700 *m/z* in positive ion polarity. For the TIMS settings, the mode was set to custom, with a starting 1/K_0_ of 0.65 V·s/cm^2^ and an ending 1/K_0_ of 1.46 V·s/cm^2^. The ramp time was set to 50 ms with a 100% duty cycle and a ramp rate of 17.80 Hz. A customized 3×16 DIA-PASEF window method was used (Supplemental Data 2).

### Computational analysis of DIA-PASEF relative quantification data in BE(2)-C cells

DIA data was analyzed with DIA-NN (35), where a spectral library search was performed against an *in silico* spectral library generated via a modified AlphaPeptDeep library generation tool (39) using a *Homo sapiens* FASTA database (downloaded 02/2024) and an in-house generated contaminant database. The fragment ion *m/z* range was set to 200-1,800, variable PTMs included were protein N-terminal methionine excision and protein N-terminal acetylation, *in silico* digestion with trypsin with cleavage at lysine and arginine for the FASTA database, a maximum of 1 missed cleavage, a peptide length between 7 to 30 amino acids, a precursor *m/z* range of 300 to 1,800, precursor charge state of 1 to 4, a fixed cysteine carbamidomethylation, a fixed mass accuracy of 15 ppm for MS2 and 20 ppm for MS1 scans, and a spectral library was generated from the DIA runs. The resulting DIA protein and peptide identifications were used for downstream parallel reaction monitoring (PRM).

### Targeted-MS (PRM-PASEF) acquisition for relative quantification in BE(2)-C cells

The resulting DIA spectral library was imported into Skyline (36) to identify peptide candidates for targeted PRM analysis. Peptides from target proteins were selected according to presence in the DIA spectral library, and a scheduled PRM method was designed using the same DIA method settings, except the scan mode was set to PRM-PASEF. An inclusion list was created from the detected peptides in Skyline and imported into the PRM-PASEF method. The resulting PRM-PASEF data files were analyzed in Skyline, where co-eluting peaks were selected for downstream quantification if the peak was within 5 minutes of an identification made during the DIA-PASEF runs and made with a minimum of 5 fragment ions. Summed integrated peak areas of the extracted fragment ion chromatograms (XIC) for 3 fragment ions per precursor were used for quantification. A minimum XIC dot-product score of 0.9 was required for each peptide for downstream quantitative comparisons. For differential detection between protein abundances, peptides were aggregated if their CV was < 20%. Fragment ion peak areas were scaled to the TIC for each run, and peptides from the same protein were averaged after row normalization for each peptide.

### Synthetic peptides for standard isotope dilution assays in BE(2-C cells

Using the spectral libraries and DIA dataset generated from the above procedure, signature peptides were selected for synthesis as light and heavy isotopologues from the following proteins: HTT, HAP40/F8A1, EXO1, FAN1, MLH1, MLH3, MSH2, MSH3, MSH6, PMS1, PMS2, GAPDH, H4, and PKM. Peptides were evaluated by summed intensities, the number of detections across different experiments/sample types, and chromatographic peak shape. If multiple signature peptide candidates were high scoring, sequences that were identical between human and mouse proteins were selected. In total, 30 target peptides and 3 control peptides (from GAPDH, H4, and PKM) were selected for synthesis by Biosynth (Supplemental Data 3). Light peptides were synthesized and purified to ≥ 95% purity and obtained as net peptide amounts by amino acid analysis. Heavy peptides were synthesized and HPLC-purified as crude peptides with either ^13^C ^15^N -Lys or ^13^C ^15^N -Arg (99% enrichment) at the C-terminus. Immediately prior to use, lyophilized peptides were resuspended in 0.1% formic acid at 1 nmol/μL and stored as single use stock aliquots. From peptide stocks, heavy-labeled and light peptides were separately combined into master mixes at equimolar concentrations (1 pmol/μl per peptide). The light peptide master mix was used to prepare serial dilutions for standard curve samples ranging from 50 fmol/μL down to 0.016 fmol/μL. The heavy peptide master mix was used to spike-in internal references at equivalent amounts (2 fmol/μL) to all blank, standard curve, and experimental samples.

### Sample preparation and DIA/PRM-PASEF acquisitions for standard isotope dilution assays in BE(2)-C cells

BE(2)-C cells were cultured, lysed, digested, desalted, and adjusted to 200 ng/μL peptide concentration, as described above in “**Sample preparation for label-free DIA-PASEF in BE**(**2**)**-C cells”**. Heavy-labeled master mixes were spiked into experimental samples. Samples (0.5 μL) were analyzed in two phases: (1) Experimental samples and the highest calibration curve points were analyzed by DIA-PASEF for spectral library generation (to assist in peak picking) and for constructing RT and IM-annotated PRM-PASEF methods (as described above) and (2) acquisition of experimental samples was repeated with PRM-PASEF scheduled acquisition, followed by acquisition of all standard curve samples including blanks. Standards and experimental samples were prepared and analyzed in biological triplicate. Skyline was used to extract fragment ion chromatograms. Peak picking was manually validated. Peak integration and calculation of light/heavy (L/H) ratios was performed in Skyline and exported to Excel and Prism for downstream analysis. Standard curves were constructed by plotting L/H ratios vs standard peptide amounts and fitting a linear fit line using 1 / y weighting. Concentration of unknowns was calculated by interpolation in Prism and expressed either versus total protein in analyzed sample (100 ng) or versus respective femtograms of GAPDH.

### Sample preparation, MS acquisition, and computational analysis of striatal tissues from Q140 and Q20 knock-in HD mouse models

Nuclear-cytoplasmic fractionation was performed in quadruplicate using the NE-PER kit (Thermofisher Scientific), as per the manufacturer’s instructions, on frozen mouse striatum for four genotype-age samples (Q20-8wk, Q140-8wk, Q20-40wk, Q140-40wk) (Jackson Laboratory, CHDI-81003005). Following biochemical fractionation, the protein yield was determined by the BCA assay. 50 μg of protein was used for trypsin digestion, as described above. Samples were generated for experimental spectral library generation by separately pooling aliquots of the cytoplasm and nuclear enriched fractions, which were then separated into 8 analytical fractions by basic reverse phase spin columns (Thermofisher Scientific). LC-MS/MS analysis (1 µL of sample containing 150 ng of peptides on column) was performed on a nanoElute 2 coupled to a timsTOF Ultra mass spectrometer (Bruker). The mobile phases were 0.1% LC-MS grade FA (Thermo Fisher Scientific) in 99.9% UHPLC-MS water (Fisher Scientific, buffer A) and 0.1% FA in 99.9% UHPLC-MS acetonitrile (Fisher Scientific, buffer B). A one column method was used with a PepSep ULTRA (25 cm x 75 μm x 1.5 µm) C18 HPLC column (Bruker) and a 10 μm emitter (Bruker) attached to a CaptiveSpray Ultra source with a column toaster set to 50°C. For analysis of pooled samples for spectral library generation, a linear 90-minute gradient from 4% to 35% buffer B at a flow rate of 150 nL/min was used for peptide separation, over which a data dependent acquisition-parallel accumulation serial fragmentation (dda-PASEF) method was employed for peptide analysis. The MS1 scan settings were set at 100 to 1,700 *m/z* in positive ion polarity. For the TIMS settings, the mode was set to custom, with a start and end 1/K_0_ of 0.65 V·s/cm^2^ and 1.46 V·s/cm^2^, respectively. The ramp time was set to 50 ms with a 100% duty cycle and a ramp rate of 17.80 Hz. For analysis of individual biological replicates, a linear 30-minute gradient from 3% to 34% buffer B at a flow rate of 200 nL/min was used for peptide separation, over which a DIA-PASEF method was employed for peptide analysis. The detector settings were the same as above, except variable width DIA windows (3 x 16 design) were specified based on optimization using the py_diAID tool (40) on dda-PASEF analysis of HeLa peptide standards (Thermo Fisher Scientific). DDA data were analyzed by FragPipe (34) (ver. 22, https://fragpipe.nesvilab.org), while DIA data were analyzed by DIA-NN (ver. 1.9, https://github.com/vdemichev/DiaNN) or Spectronaut ver. 19 (Biognosys). Results were exported to Excel for downstream processing and analysis.

### MMR interactome in BE(2)-C cells using thermal proximity coaggregation (TPCA) assays and Tapioca scoring

A 5 temperature TPCA (37°C, 40.7°C, 44.6°C, 52.8°C, and 55.3°C) and quantitative DIA MS analysis was performed in HTT KO and SCR control BE(2)-C cells, as previously described (41). Melting curve data was collected in triplicate per condition using label-free DIA analysis on a Bruker nanoElute2 HPLC and timsTOF Ultra. To obtain MMR interactions, the data were analyzed by FragPipe’s (ver. 24) diatracer module (33). Temperature curves were assembled, pre-processed, and filtered to retain curves that had ≥ 4 points in both KO and SCR conditions in ≥ 2 replicates. Pairwise interactions were scored by Tapioca using its full model prediction (42). MMR-MMR and MMR-non-MMR pairwise interactions were retained with Tapioca scores ≥ 0.75 and ≥ 0.80, respectively. Cytoscape (ver. 3.10.4) was used to visualize literature interactions for the MMR interactome using the STRING database, or to directly visualize Tapioca predicted interactions.

## Results

### Selection of signature proteins in DNA repair and handling pathways for development of targeted-MS assays

Our goal was to develop sensitive and selective quantification assays using targeted-MS to provide a unified method to quantify proteins involved in DNA repair and handling pathways (Figure 1A-B). The selection of high priority targets for assay development was based on pathway members that are implicated in modulating CAG repeat expansion in Huntington’s disease (2, 9, 10), such as mismatch repair proteins (Figure 1A). Moreover, accumulating evidence from genome-wide association (GWA) and HTT protein interaction studies points to a broader role of factors that impact disease onset and progression, including DNA ligases (LIG1), FAN1 exonuclease, polymerases (POLD1), helicases (ERCC3), and even transcriptional regulators (TCERG1) (3, 43–46). In addition, functionally related proteins, e.g., other POL and LIG proteins, were included in the assay development so secondary or compensatory changes in the DNA R/H landscape could be monitored. In total, 43 proteins (Figure 1B and Supplemental Data 4) were included for assay development using platform-optimized, MS-based quantification workflows (Figure 1C).

**Figure 1.**
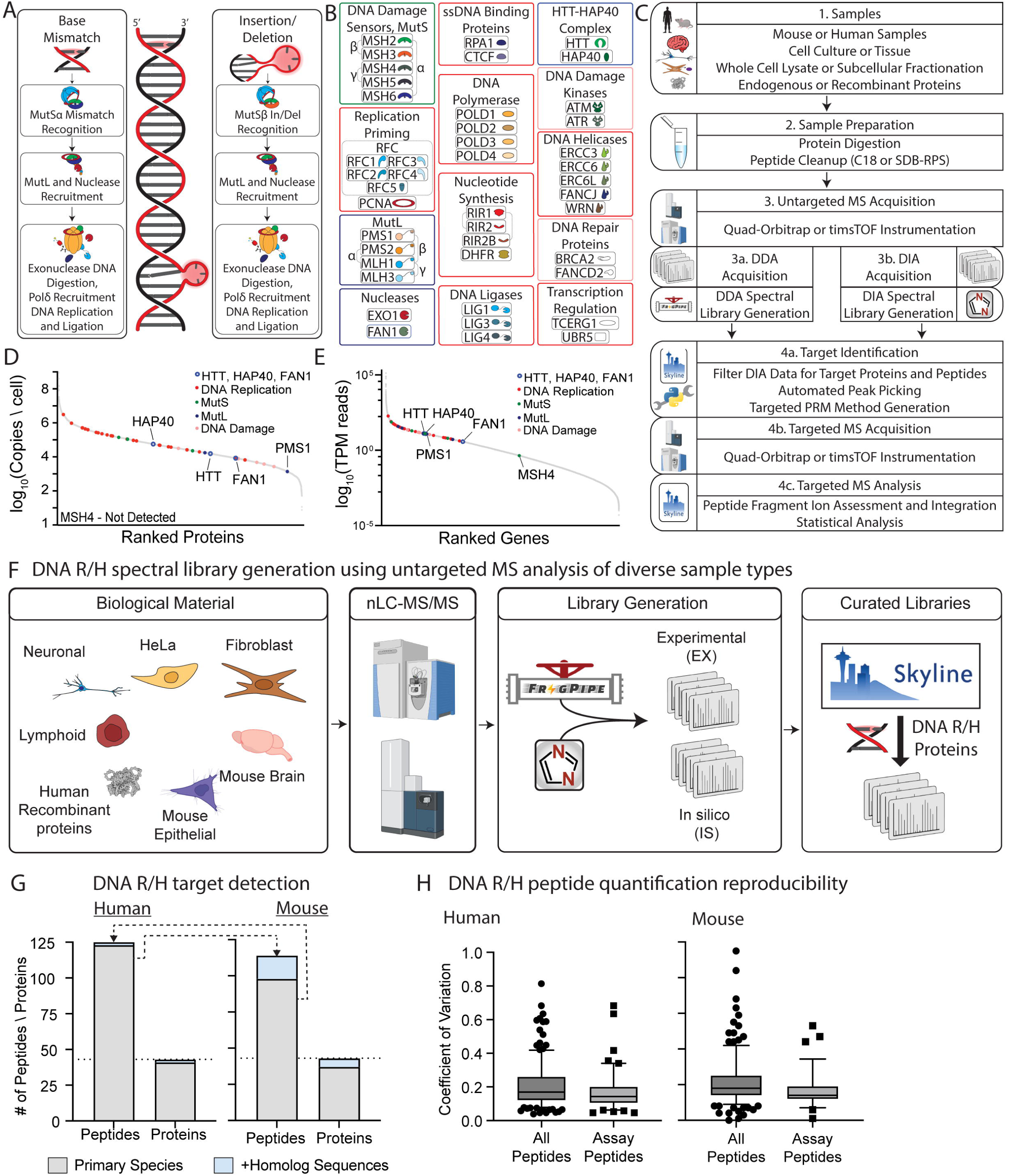
Overview of human and mouse PRM assay construction and properties. (A) Simplified schematic of the DNA repair pathway for base mismatch and insertion/deletions event. Mismatch repair protein complexes and accessory proteins are highlighted. (B) Targeted MS assays were developed using these DNA repair and handling proteins, which are shown assigned to general molecular function classes. (C) Unified workflow for developing targeted-MS assays suitable for timsTOF, quadrupole-Orbitrap, or triple-quadrupole platforms, which outlines a stepwise DDA/DIA-to-PRM strategy. (D & E) Representative abundance ranking of proteins (D) and RNA (E) from HeLa and neuroblastoma cells, respectively, with key DNA R/H targets highlighted. (F) Detailed workflow for untargeted MS acquisition, illustrating biological samples that were characterized for generation and curation of DNA R/H spectral libraries. (G) Number of peptides and proteins manually curated from spectral libraries generated from (F) for human (left) and mouse (right) samples. Horizontal dotted line shows complete assay coverage (proteins). (H) Coefficient of variance from DIA analysis of all or selected DNA R/H peptides in human BE(2)-C cells and mouse striatum.

We established two workflows for constructing targeted-MS assays (Figure 1C). One workflow was tailored for liquid chromatography-based platforms coupled to widely available Orbitrap-based or triple quadrupole mass spectrometers. Historically, these platforms are used to develop gold-standard MRM and PRM targeted-MS methods (28, 47, 48). Yet, given advances in timsTOF and Astral MS platforms, this study evaluated key aspects of assay development for these high-performance platforms, including spectral library generation, MS acquisition method, and computational pipelines. We leveraged their increased sensitivity and high scan rates (up to ∼300 precursors/sec) to establish a data independent acquisition (DIA)-driven PRM approach that offers faster development time, increased assay flexibility, and inter-lab adaptability. The basis for these assays is the reproducible detection of proteolytic signature peptides that serve as surrogates for protein abundances. For PRM assays developed on ultra-fast scanning instruments, 500-1000 peptides can be routinely supported in a single injection per sample (49). In contrast, for targeted assays on Q-Exactive (QE) platforms, a reasonable scope is ∼100-150 monitored precursors per injection (for typical 60 min LC gradient lengths).

### Generation of spectral libraries from human and mouse model systems drives signature peptide selection for Orbitrap-based assays

When considering the protein abundances of the DNA R/H targets of interest (24), we noticed that these span most of the detectable range, ∼4-orders of magnitude (Figure 1D). Therefore, compared to the detection of these targets at the RNA level (Figure 1E) (23), protein quantification presents a greater challenge. Accordingly, we constructed experimental spectral libraries of tryptic peptides generated from diverse human and mouse cells/tissues, which were analyzed by 1D and 2D reverse phase nanoLC combined with data-dependent and independent acquisition MS/MS approaches (Figure 1F). These different samples and acquisition strategies increased our chances to detect challenging targets and established the subset of reproducible signature peptides. This was an essential step for establishing QE-based assays, where our goal was to quantify all targets in a single analysis with an average of ∼3 peptides/protein. Of the 43 DNA R/H targets, 41 and 37 were detected for human and mouse libraries, respectively (Figure 1G, gray bars). Using computational “transfer” of homologous sequences between species-specific libraries (Supplemental Data 5), the theoretical target coverage was complete, reaching 43 proteins for both libraries (Figure 1G, gray+blue bar).

Based on the peptides detected for the target proteins, we calculated observability, intensity rankings, and coefficient of variances across different datasets (Supplemental Data 5). Using these metrics, 125 and 114 peptides were selected as signature peptides for the 43 target proteins in the human and mouse libraries, respectively. Notably, the signature peptides were measured with lower CV (< 25%) compared to all detectable targeted peptides (Figure 1H). Overall, most target proteins were represented by 3 peptides, except for MSH4 and MSH5, which had single peptides. We also selected control peptides from enolase, histone H4, and PKM for additional normalization purposes. To accelerate the setup of DNA R/H assays in other laboratories with different instrument configurations, we assembled the assay targets and respective peptides in species-specific Skyline template documents that contains predicted retention time and MS/MS fragmentation spectra (PanoromaWeb project; https://panoramaweb.org/DNARH_PRM.url), which can be used to generate platform-specific targeted-MS instrument methods and analyze future quantitative datasets.

### Optimizing a computational pipeline for DIA-driven targeted MS workflows

Software developments have the potential to provide immediate gains without the requirement to perform new experiments. To evaluate the benefit of different DIA computational pipelines for analysis of brain tissue from HD mouse models, we compared performance metrics between two common software platforms, DIA-NN (DN) and Spectronaut (SN), used with the above described experimental (EX) library or an in silico predicted (IS) library (Figure 2A). For this analysis, we selected a mouse HD model system, as detection of the DNA R/H targets in mouse samples proved more challenging (Figure 1G). Specifically, striatum brain tissues were dissected from mice expressing mutant huntingtin (HTT-Q140) or wild type HTT (Q20) at 8 and 40 weeks of age (Figure 2A). Dissected striata were separated into nuclear and cytoplasmic-enriched fractions and pooled for spectral library generation. The timsTOF DDA-PASEF analysis of nuclear and cytoplasmic-enriched proteomes was used to generate the EX library using FragPipe. Overall, the EX library contained ∼9600 proteins and ∼175k precursors (filtered at 1% FDR).

**Figure 2.**
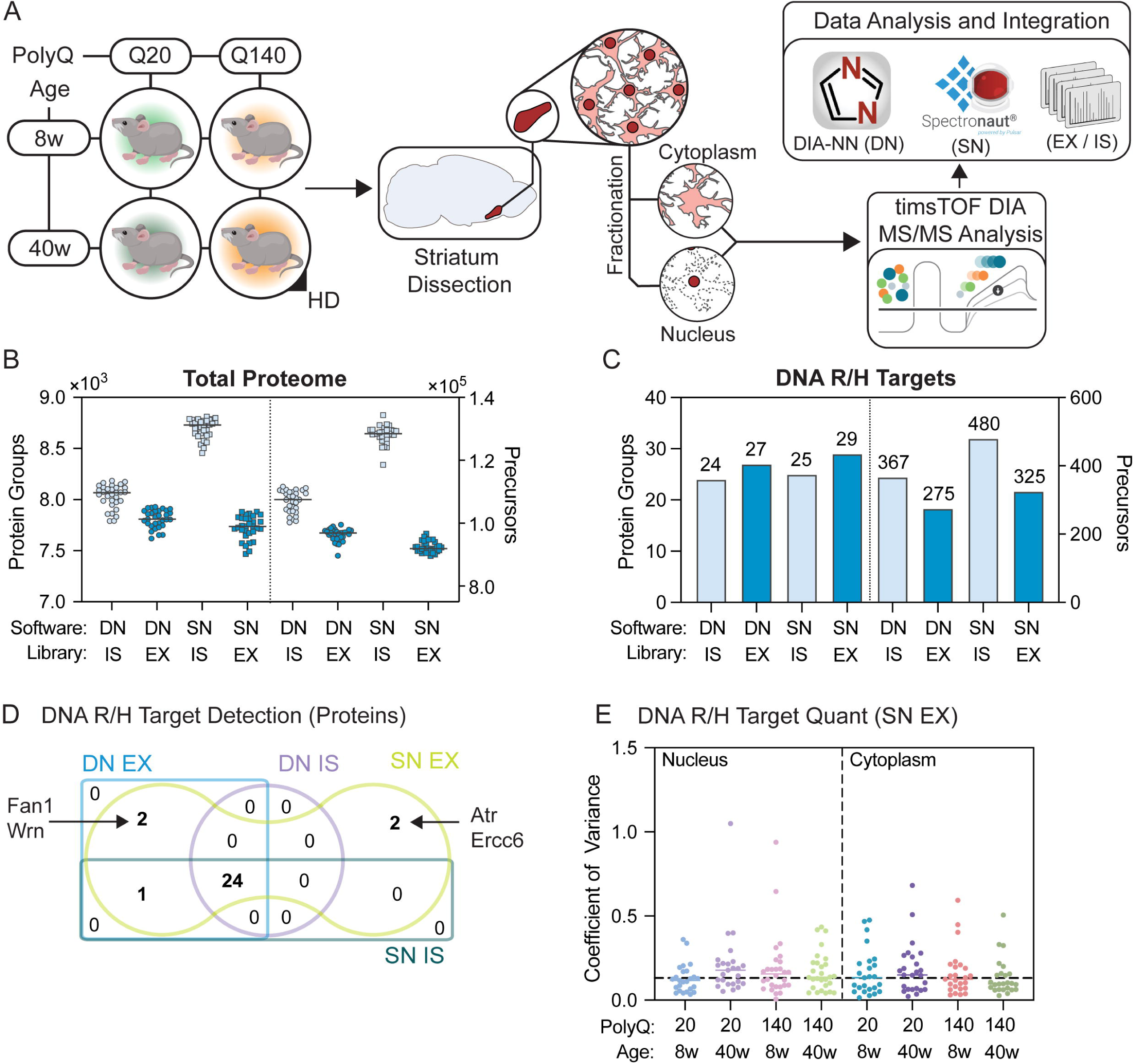
Benchmarking and optimizing DNA R/H assay computational pipelines in HD mouse models. (A) Striata from HD and control knock-in mouse models at 8 and 40 weeks of age (n = 4 replicates) were subjected to subcellular fractionation of nuclear- and cytoplasmic-enriched fractions, analyzed by DIA-PASEF and processed by DIA-NN v1.9 (DN) or Spectronaut v19 (SN) computational pipelines. (B) Number of protein groups and precursors identified by DN or SN using either a mouse in silico predicted (IS) or experiment-specific (EX) spectral library. Distributions show individual results from samples generated in (A) (n = 32). (C) Protein groups and precursors identified for DNA R/H targets. Data is reported as the union of all samples from (A). (D) Overlap of proteins identified in (C) among four computational workflows. (E) Coefficient of variance (CV) distributions for the best performing workflow, SN-EX, separated by mouse genotypes (Q20 & Q140), age (8 and 40 weeks), and subcellular fraction. Each point represents the CV for one target protein.

The DIA-PASEF identification performance was next compared for non-pooled striatum samples (Figure 2A) using either IS or EX libraries in DIA-NN and Spectronaut DIA analysis software. The best overall performance at the total proteome level was with Spectronaut using an IS library (SN-IS), identifying 33% more precursor ions and 11% more protein groups than DIA-NN using the mouse striatum EX library (Figure 2B). Independent of the software, the IS library outperformed the EX library (Figure 2B, grey versus green circles). However, this pattern at the proteome level did not hold for detection of DNA R/H proteins, where EX library searches had greater target detection (Figure 2C), with Spectronaut slightly outperforming DIA-NN (29 vs 27 proteins). The EX library searches produced lower precursor numbers per protein (Figure 2C). Notably, FAN1 was identified when searched against the EX but not IS library (Figure 2D). While the greater protein-level sensitivity of the EX library was at the expense of more missing quantitative values, this trade-off was acceptable given the detection of four additional targets while still achieving median variances of < 15% (Figure 2E).

### Mutant HTT has altered levels and subcellular distribution, but does not alter DNA R/H protein abundances in a Q140 knock-in mouse model

Using our optimized computational pipeline (Figure 2), we investigated if DNA R/H targets in an HD Q140 mouse model (50, 51) were differentially regulated by mutant HTT, and whether the changes were dependent on age or subcellular localization. In the nuclear-associated fraction of Q140 mice, we observed that HTT and HAP40, a key structural and functional partner of HTT (52, 53), were reduced compared to the control (Q20) mice at both 8 and 40 weeks (Figure 3A & 3B), consistent with our and previous proteome studies of these knock-in HD mouse models (54, 55). The downregulation of HTT was significant at both 8- and 40-wks, while HAP40 was only significant at 40wks. For the quantified DNA R/H targets, none were dramatically altered by Q140 HD pathogenesis, but several trended lower (RRM2B, RPA1, MSH2, and ERCC3). Interestingly, the abundances of RRM2B and MSH2 were increased due to aging in Q20 and Q140 genotypes, respectively (Figure 3C and 3D). As the measured DNA R/H target abundances were not notably changed in HD mice, we examined whether mHTT impacts the targets’ subcellular distribution. The subcellular distribution of the core DNA R/H targets did not significantly change in HD mice. However, for HTT and HAP40, both showed a shift in localization from the cytoplasmic- to nuclear-associated fraction at 40wks, though the bulk of each protein remained preferentially cytoplasmic (Figure 3E). Previous studies of HD mouse models have shown increasing accumulation of nuclear HTT, but this reflected largely N-terminal HTT fragments that readily form fibrillar insoluble aggregates (56–58). Therefore, this observation provides evidence that soluble, full-length mutant HTT shifts toward nuclear-associated compartments during HD progression.

**Figure 3.**
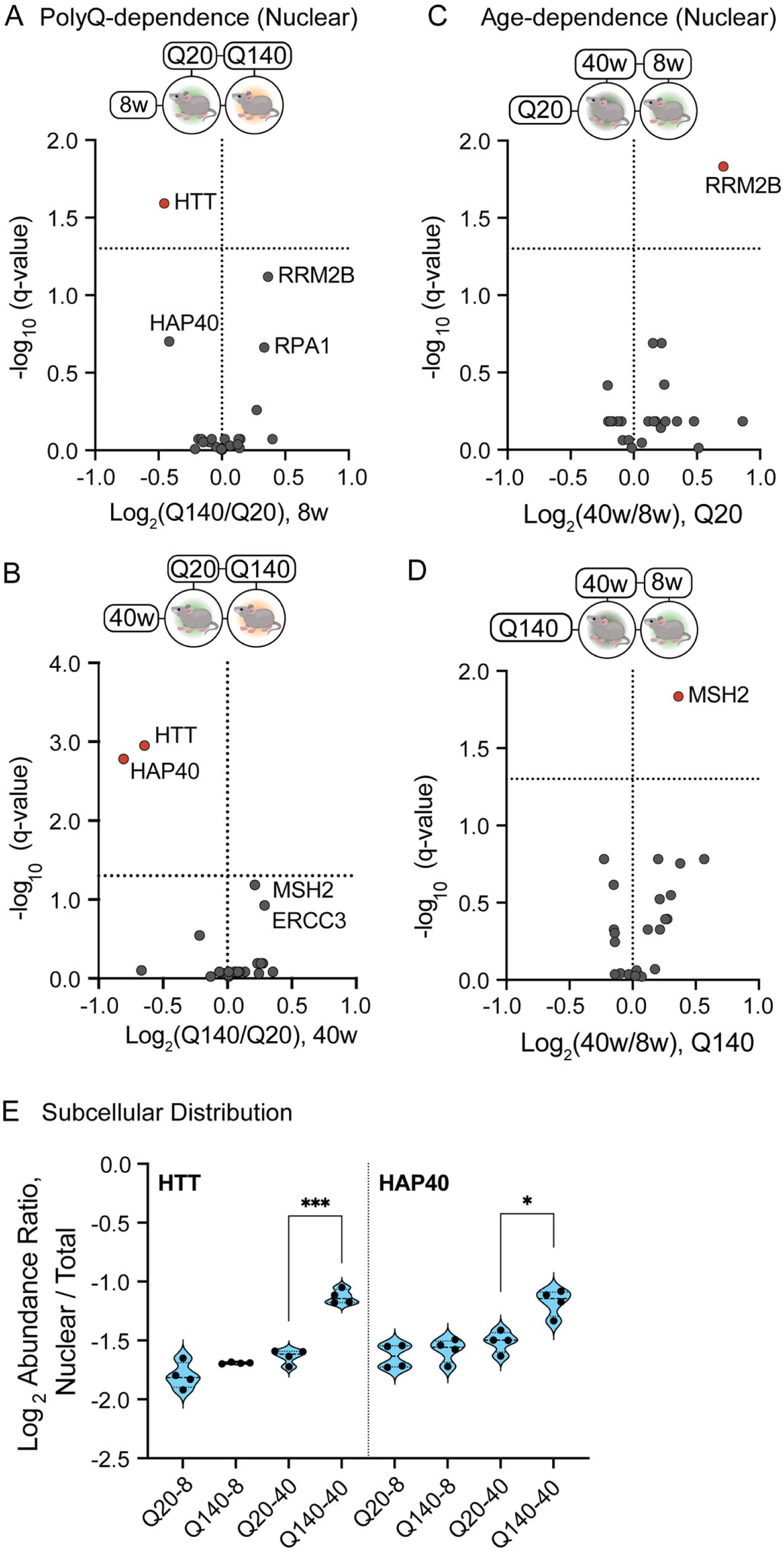
Evaluating the effect of mutant HTT and disease progression on DNA R/H protein abundance in the striatum of HD KI mouse models (see. Fig. 2A). (A - D) Differential abundance analysis of DNA R/H targets in nuclear-enriched striatal fractions as a function of expanded polyQ (Q140 vs Q20) at 8wk (A) and 40 wk (B), and age in Q20 (C) and Q140 (D) mice. Volcano plots illustrate the magnitude (expressed as log2 fold-change) and significance (q-value < 0.05) of the relative abundances. Significant detections are indicated as red colored symbols. (E) Change in nuclear vs total nuclear+cytoplasm abundance fraction across genotype-age conditions for HTT and HAP40. Significance testing compared means by t-tests (***p < 0.001, *p < 0.05, n = 4).

### Development of an absolute quantification assay for DNA R/H proteins in an HTT knock-out human neuroblastoma cell model

Having confirmed our ability to detect the DNA R/H targets in mouse model systems, we next turned our attention to human cells. While mouse models recapitulate many features of the disease in humans, for example, CAG expansion and transcriptional dysregulation, other facets, such as the near-complete striatal degeneration and few intranuclear inclusions are not well represented (59). Therefore, characterization of HD in human cell models can reveal unique features that may facilitate future potential translational research applications. Our assessment of prior proteome datasets suggested that the challenges in detecting these proteins may be in part linked to their range of protein abundances in different cell or tissue types (Figure 1D & 1E). To date, no multiplexed assay is available to rigorously quantify DNA R/H proteins. Therefore, we sought to obtain a clearer picture of their abundances through absolute quantification by mass spectrometry. As a first step in the development of absolute quantification assays, light and heavy isotopic standard peptides were synthesized for 11 of the DNA R/H targets that were of prioritized therapeutic interest: HTT, HAP40, EXO1, FAN1, MLH1, MLH3, MSH2, MSH3, MSH6, PMS1, and PMS2 (Supplemental Data 3). Our existing spectral libraries and DIA datasets were used to determine the top 2-4 candidate sequences for each protein. We evaluated the peptides’ summed intensities, the number of detections across different experiments/sample types, and chromatographic peak shape. When multiple signature peptide candidates were high scoring, we selected sequences that were either identical or highly similar in the mouse proteins. In total, 30 peptide sequences were selected and synthesized as light and heavy isotopologues. Light peptides were synthesized to ≥ 95% purity and used to construct the standard concentration curves. Heavy peptides were synthesized and HPLC-purified as crude peptides with either ^13^C_6_^15^N_2_-Lys or ^13^C_6_^15^N_4_-Arg at the C-terminus. These peptides were used as internal spike-in standards added at equivalent amounts to all blank, standard curve, and experimental samples (Figure 4A). We first performed in-house validation of synthetic peptides and constructed response curves in HeLa cells. Specifically, we observed that peptide intensities of standards measured in a matrix of 10 ng/μL HeLa lysates were similar to neat standards (standards alone) (Figure 4B). Nonetheless, any differences in ion suppression between light synthetic peptides in calibration curves and endogenous peptides in experimental samples are corrected by using isotope (L/H) ratios. Hence, we pursued assay development using external standard curves, which conserves experimental samples that may be in limited supply. We constructed standard curves with standard (light) peptide concentrations covering 0.016 fmol/μL to 50 fmol/μL and an internal standard (heavy) peptide concentration of 2 fmol/μL to minimize detection of the low amount of light impurity (∼0.5 %) (Figure 4A).

**Figure 4.**
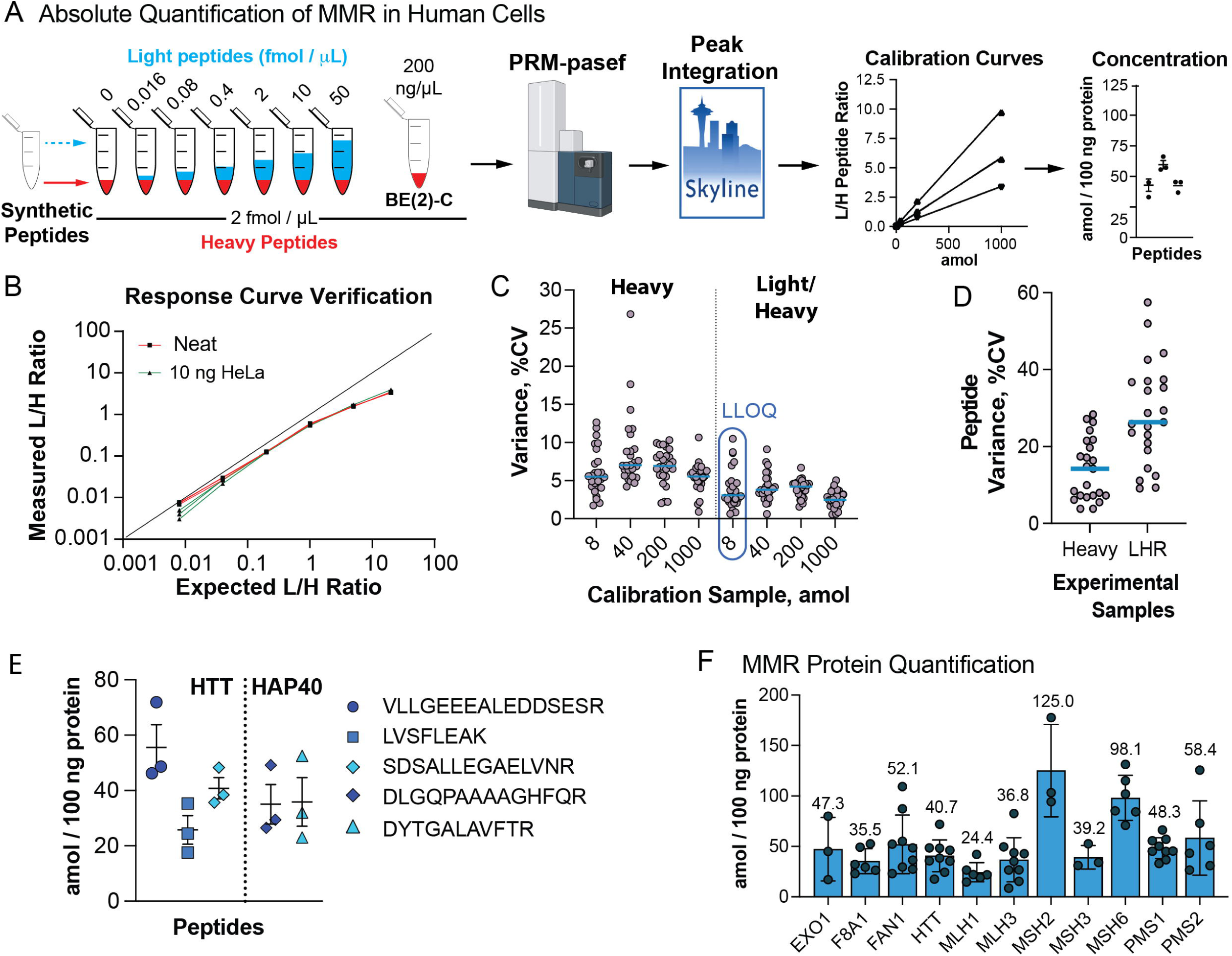
Absolute quantification of DNA R/H proteins in wild-type BE(2)-C neuroblastoma cells. (A) Standard isotope dilution experiments using PRM-PASEF acquisition were performed with experimental samples from WT BE(2)-C tryptic digests with calibration curves. (B) Representative peptide light-to-heavy (L/H) ratio response curve plotted for expected versus measured ratios in two backgrounds: neat and HeLa peptide matrices. (C) Coefficient of variance (%CV) distributions for 27 synthetic peptides at calibration points of 8, 40, 200, 1000 amol. CV is plotted for heavy peak areas and light/heavy ratios. The calibration point reflecting an empirically defined lower limit of quantification is indicated. (D) Similar to (C) except % CV was plotted for heavy peptide peaks areas and Light / Heavy ratios (LHR). (E) Concentration of the three HTT and two F8A1 peptides. (F) Target protein concentrations relative to total protein. Individual data points are shown for each peptide and replicate. Mean +/- SD (n = 3 biological replicates).

We next measured absolute protein concentrations of the 11 MMR-associated proteins in BE(2)-C neuroblastoma cells, a human cell type previously used in HD studies (54, 60, 61). Tryptic digests from these samples were prepared as above, in triplicate, and heavy labeled internal standard peptides were spiked in (Figure 4A). We applied our DIA-driven PRM workflow (Figure 1C) to maximize quantitative performance for standard curve and experimental samples. Following PRM-PASEF analysis, the fragment ion chromatograms were extracted and integrated using Skyline (36). Evaluation of calibration curves showed saturation at 50 fmol/μL. After discarding the saturated points, 27 out of 30 DNA R/H peptides showed linear fits with r^2^ ≥ 0.99 (Supplemental Data 6); a single peptide from EXO1, MSH2, and MSH3 showed lower replicate reproducibility (r^2^ < 0.95) Also, one synthetic peptides from FAN1 (TNLTPGQSDSAK) had poor reproducibility. Together these four peptides were excluded from downstream calculations of protein concentration. We evaluated measurement reproducibility and lower limit of quantification (LLOQ) for the standard curve peptides. The measurement variance of heavy peptide peak areas and L/H ratios was appropriate for PRM assays, with the majority below 15% coefficient of variance (CV) (Figure 4C, *Heavy*). The 40 amol calibration point had slightly higher CV due to systematically lower signal in one replicate but was corrected for after calculating L/H ratios, which lowered CV across all points (Figure 4C, *Light/Heavy*). These measurements also allowed us to establish the peptides’ theoretical lower limit of quantification (LLOQ). A commonly accepted LLOQ definition is the lowest standard curve point where peptide L/H ratios are measured with a CV of < 20% (62, 63). At the lowest concentration point of 8 amol on-column, all L/H ratios were measured with % CV below this threshold (< 11%), with a median of 3% CV (Figure 4C). This shows the excellent reproducibility of peptide measurement for the standard curves.

For the experimental BE(2)-C tryptic digests, measurement variance increased compared to the standard curves but was still within acceptable ranges, with a median measurement variance of ∼18% and ∼25% for the heavy peak areas and L/H ratios (LHR), respectively (Figure 4D). Increased LHR variance compared to heavy peak areas was expected in the experimental samples since measurement of endogenous (light) peptides is subject to biological and sample preparation variances. The concentrations of endogenous peptides in the experimental samples were calculated using linear fits of the calibration curves (Supplemental Data 6). Overall, most peptides from the same proteins were consistent, and all peptides and their replicate measurements were averaged. For example, 2 of the 3 peptides for HTT showed similar concentrations, and the two peptides from HAP40/F8A1 were highly consistent (Figure 4E).

Comparing the calculated protein concentrations in BE(2)-C cells, most MMR proteins fall within a similar abundance range. MLH1 had the lowest measured concentration at 24.4 amol/100 ng of protein, while MSH2 and MSH6 had the highest concentrations (Figure 4F). While knowledge of absolute protein measurements is scarce, especially for MMR proteins, Beck and colleagues (64) performed absolute quantification of ∼50 reference proteins in a human osteosarcoma (U2OS) cell line, which were used to indirectly estimate the concentrations of all other proteins measured by label-free quantification. They proposed that the dynamic range of abundances within the global proteome likely spans from 20 to 20,000,000 copies per cell (64). To put our measurements into context, we selected for reference a hypothetical medium abundance protein at 200,000 copies per cell; the equivalent of 166 amol/100 ng (assuming 0.2 ng total protein per cell). All of the DNA R/H proteins that we quantified in BE(2)-C cells were below this concentration (Figure 4F). For example, in these cells, the MSH2 concentration was modestly higher (125 vs. 82 amol/100 ng) while MSH6 was 3-fold higher (98 vs. 32 amol/100 ng) compared to U2OS cells. HTT and HAP40 concentrations were ∼30 times higher in BE(2)-C compared to U2OS cells. More broadly, literature reports of HTT concentration in human samples are limited. Absolute HTT quantities have been estimated at 10 amol/100 ng in HeLa cells using a mass spectrometry-based “proteomic ruler” technique (65) and at 15 amol/100 ng in HD-derived lymphoblasts using an antibody-based MSD assay (66). The concentrations we measured in cells were 3 – 4 times higher. Overall, to our knowledge, our reported concentrations provide the first absolute quantitative measurement of mismatch repair proteins in human cells and provides a framework for future implementation of this assay in clinically relevant human derived cells, e.g., lymphoblasts, fibroblasts, IPSC-derived neurons.

### Relative quantification of DNA R/H proteins in a human neuroblastoma HTT KO model

While absolute protein concentrations are useful for certain scientific and clinical goals, a more common need is relative quantification of the same protein(s) in different biological and disease states. This is powerful when paired with DIA or targeted-MS approaches. Our DIA-based quantification of DNA R/H proteins from mouse brain on the timsTOF platform (Figure 2) hinted at similar quantitative performance to a PRM-based assay. Therefore, we proceeded to systematically evaluate the ability of these acquisition strategies (DIA vs PRM) to perform differential detection of DNA R/H proteins in human cells on the timsTOF platform. While HTT pathobiology clearly intersects with mismatch repair processes (2, 7, 15), an open question is if the biology of the wild-type protein modulates MMR or DNA R/H pathways. Towards this goal, a KO of HTT was generated in BE(2)-C cells using CRISPR technologies. We observed HTT KO in ∼40-50% of cells, consistent with a 40% reduction in HTT protein at the bulk proteome level (Figure 5A). This result mimics the magnitude of reduction that has been achieved in therapeutic HTT lowering clinical trials.

**Figure 5.**
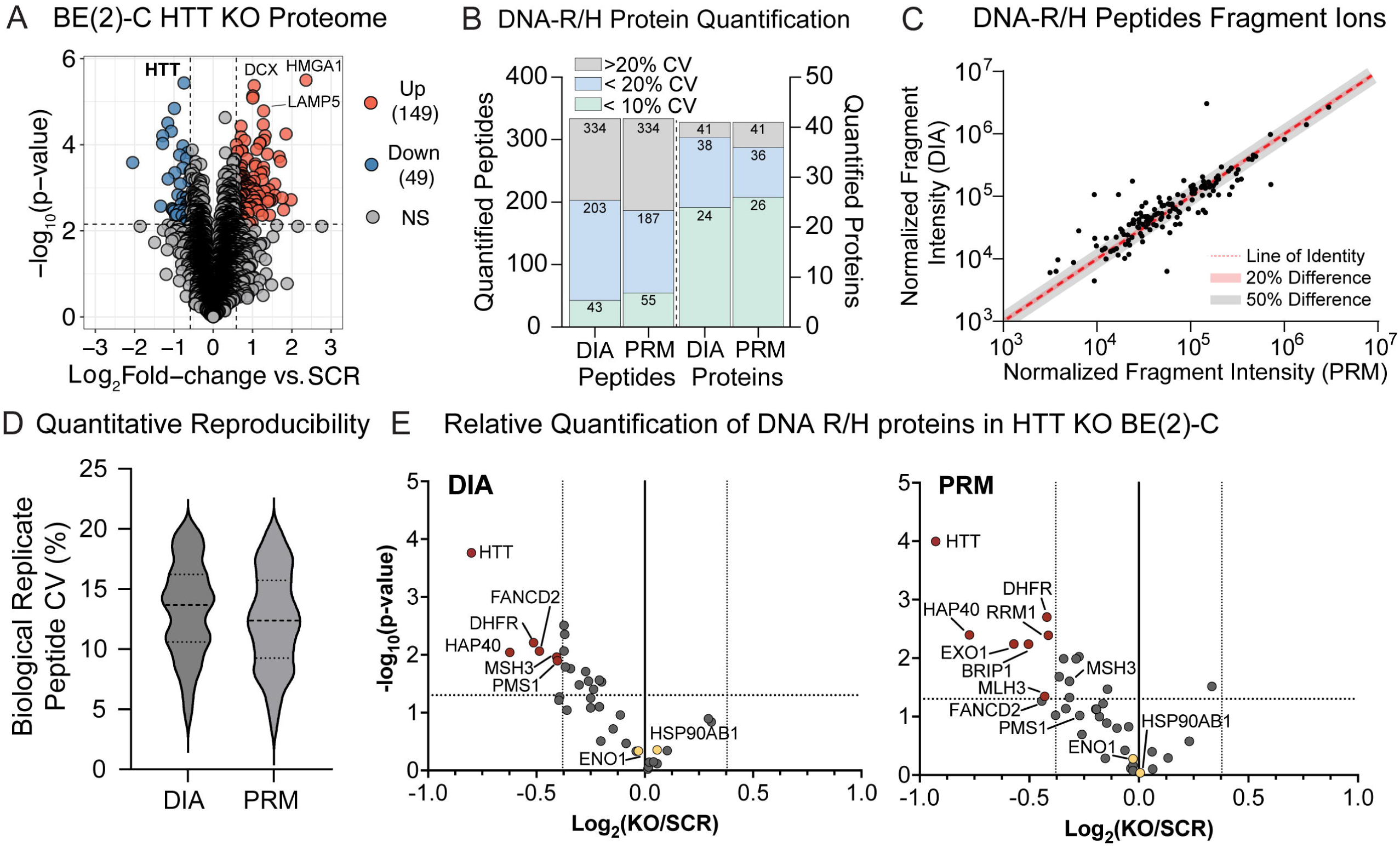
Relative quantification of DNA R/H proteins in HTT KO BE(2)-C neuroblastoma cells. (A) Differential proteome abundance analysis due to HTT KO vs control (SCR) in BE(2)-C cells (n = 3 replicates). Vertical and horizontal dashed lines indicate thresholds for significance, FC ±1.5-fold and p-value < 0.05, respectively. (B) Number of total precursors and proteins that were quantified in BE(2)-C SCR cells using DIA and PRM, stratified by coefficient of variation (CV). (C) Fragment ion intensities of shared peptides from DNA R/H proteins measured by both DIA or PRM. (D) Distribution of peptide CV measured by DIA-PASEF or PRM-PASEF analysis of BE(2)-C SCR. (E) Differential detection of DNA R/H proteins using DIA (left) or PRM (right) quantification was visualized by Volcano plots. Protein abundances were calculated from the average precursor abundances (n = 3 replicates) with < 20% CV within each quantification approach. Red and yellow-filled circles are differential targets and endogenous proteins used for normalization, respectively. Vertical and horizontal dashed lines indicate thresholds for significance, FC ±1.3-fold and p-value < 0.05, respectively.

Using this human cell model, we compared the detection capability of DIA and PRM for the DNA R/H targets from BE(2)-C cells. For a validation comparison of DIA- and PRM-based quantification, the same peptides and fragment ion transitions were manually selected and integrated in Skyline for both acquisition strategies. Comparing DIA and PRM acquisitions, the total number of quantified DNA R/H targets was identical (Figure 5B) and fragment ion intensities were largely similar (Figure 5C). However, stratifying peptide and protein quantification by measurement variance, revealed an increase of ∼20% in the number of precursors that were measured by PRM at < 10% CV compared to DIA (Figure 5B). Moreover, comparison of the relative CV for the same peptide fragment ions showed slightly lower CV for PRM (Figure 5D). Overall, we found that PRM enabled slightly higher dynamic range and reduced measurement variance, but similar detection sensitivity to DIA.

We next investigated if the analytical performance benefit of PRM would translate to superior differential detection of DNA R/H targets in the context of HTT KO. For this comparison, precursors measured with < 20% CV were retained, averaged to the protein level, and relative abundance expressed as KO versus SCR control. In HTT KO cells, PRM and DIA showed statistically significant reductions in HTT, HAP40/F8A1 and DHFR (Figure 5E, *left vs right*). PRM-based quantification provided an additional 4 differential targets (EXO1, BRIP1, RRM1, and MLH3) (Figure 5E, *right*). EXO1 was a differential target only by PRM because this acquisition improved measurement of its peptides to < 20% CV compared to DIA. While generally protein relative abundances from PRM were larger than DIA for the same target, the ratio for MSH3 was lower and below threshold in PRM, though it was still statistically significant (Figure 5E, *right*). FAN1 was only quantified by one peptide with a CV of > 25% and was excluded, which showed an increasing trend with HTT KO. This contrasts with other Fanconi anemia pathway members, BRIP1 and FANCD2 which were quantified by 3 and 4 precursors, respectively, and were significantly downregulated. While the relative merits of DIA and PRM acquisition for relative quantification will be discussed below, from an analytical perspective, the simplicity and flexibility of DIA quantification is attractive. It acquires data for all quantifiable proteins, which can be computationally mined for additional targets and hypothesis generation. In contrast, PRM only acquires data for the pre-defined targets, but it provides increased reproducibility and potentially sensitivity for differential detection. PRM method development has additional resource investment, but once the primary PRM assay has been established, new peptide targets can be theoretically added through in silico prediction of RT and IM parameters.

### HTT knock-out alters the protein interactomes of MutS and MutL mismatch repair complexes

Our relative quantification studies demonstrated that HTT lowering in human neuroblastoma cells impacts the levels of only a few MutS and MutL mismatch repair complex (MMR) components (Figure 5E). Yet, HTT loss/knock-out has been shown to have broad effects at various omics levels in mouse embryonic fibroblasts (67), and more recently in human SH-SY5Y neuroblastoma cells (61) and stem cell-derived cortical neurons (26). Given these systems level effects resulting from HTT loss, it is possible that MMR complexes are impacted through their functional protein interactions. We explored this hypothesis through global protein interaction analysis of HTT KO and wild-type BE(2)-C cells using thermal proximity coaggregation assays paired with mass spectrometry (TPCA-MS) (68–70)(Figure 6A). Overall, we assembled melting curves for nearly 7,000 proteins in each condition, 95% of which were quantified in at least two biological replicates (Figure 6B). Among these reproducible curves, we had high representation of proteins in DNA R/H pathways (34 proteins) and specifically mismatch repair proteins (7 proteins) (Figure 6B). This provided coverage across all core MMR complexes (MutSα, MutSβ, MutLα, and MutLβ), except for MutLγ as MLH3 was not quantified. Protein interactions were predicted using Tapioca, a machine learning tool that we previously developed for *de novo* PPI prediction by integrating information from experimental melting curves, protein properties, and known functional networks (42). Across the BE(2)-C proteome, nearly 23, 000 binary PPIs were predicted (Tapioca score > 0.8) (Figure 6C). Among these, we observed all known heterodimeric MMR interactions, MSH2-MSH3 and MSH2-MSH6 for MutS complexes, and MLH1-PMS2 and MLH1-PMS1 for MutL complexes. We also predicted the known interaction of exonuclease 1 (EXO1) with MutL via its interaction with MLH1, and sub-threshold interactions (Tapioca score 0.75 – 0.80) between MutL and MutS complexes (MLH1-MSH2, MLH1-MSH3, and PMS2-MSH6), consistent with the existence of a transient ternary complex with DNA (71, 72). These results demonstrate that TPCA-MS effectively captures the topology of MMR complexes in BE(2)-C cells, supporting further exploration of *de novo* predicted MMR interactions and their relevance to HTT biology.

**Figure 6.**
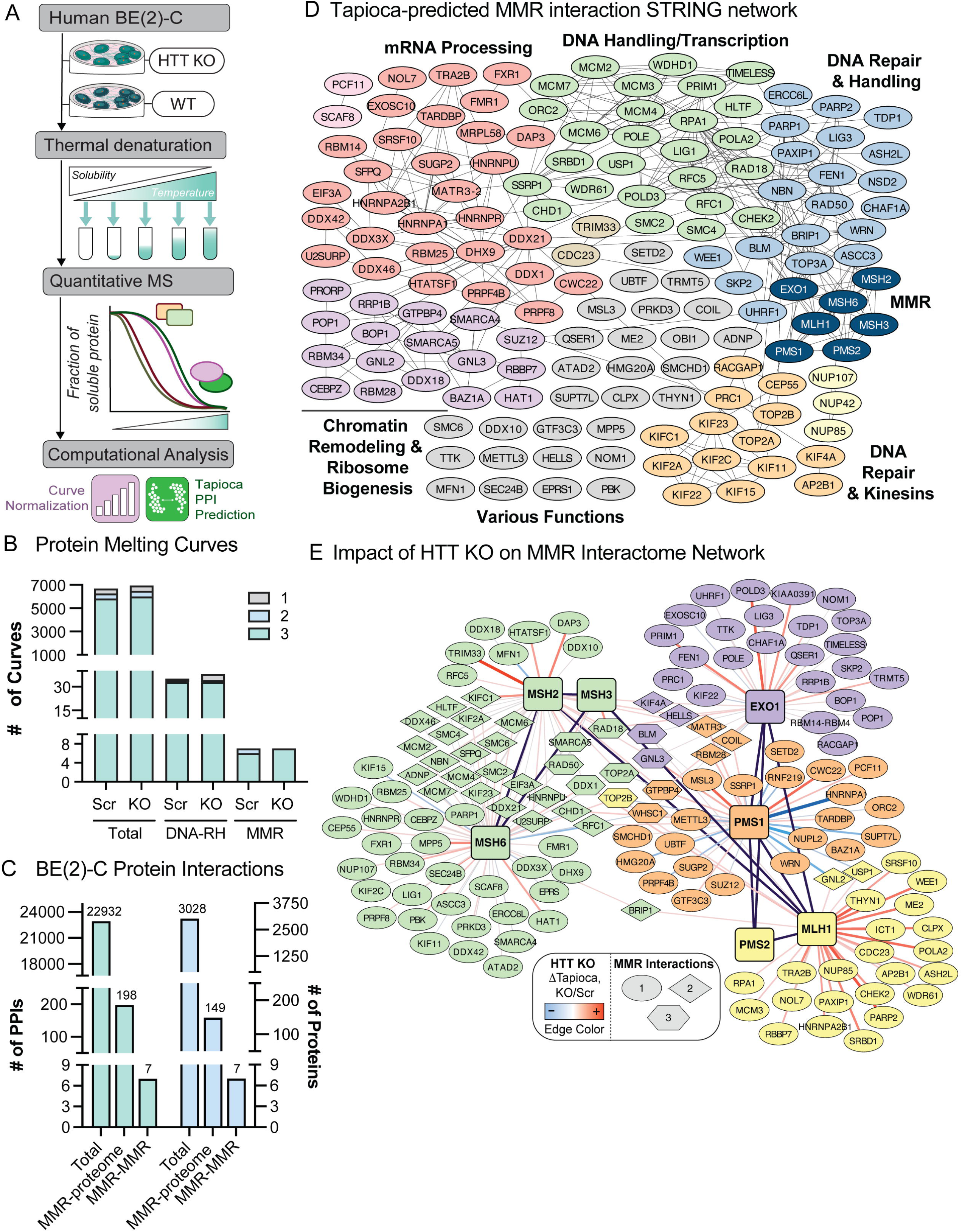
HTT KO modulates MMR protein networks. (A) Experimental workflow to characterize MMR protein interactions in HTT KO BE(2)-C cells using thermal proximity coaggregation-MS assays (B) Number of assembled melting curves for all proteins (Total), DNA repair and handling (DNA-RH), and mismatch repair (MMR) proteins. Stacked bars show the number of curves that were assembled in 3, 2, and 1 replicate for control (Scr) and HTT KO conditions. (C) Number of predicted binary protein interactions (PPIs) and respective number of proteins from Tapioca analysis of melting curves for all proteins (Total), involving MMR proteins (MMR-proteome), or between MMR proteins (MMR-MMR). (D) STRING physical interaction network of Tapioca-predicted interactions with MMR proteins. (E) Tapioca-predicted MMR network showing effect of HTT KO (edge color). Seven seed proteins (from MutS and MutL complexes and EXO1) are predicted to form interactions with 142 cellular proteins (Tapioca score > 0.8 in at least one condition).

Beyond MMR protein complex interactions, 142 interactions were predicted with at least one of 7 MMR seed proteins (six MutS/MutL subunits plus EXO1). Assembly of a physical interaction network among these proteins using the STRING database resulted in an interconnected network that bridged MMR proteins to essential proteins that facilitate DNA repair and handling during DNA damage (Figure 6D, blue, tan, and green nodes) and downstream transcriptional components that contribute to DNA damage resolution (Figure 6D, green nodes). TPCA also captured associations with proteins classically annotated to RNA binding/processing, such as DEAD-Box RNA helicases (DDX21, DDX3X), RNA-binding proteins (FXR1, FMR1, RBM14, RBM25, and HNRNPs), as well as components of chromatin remodeling complexes ISWI, PRC2, BAF, and NuRD (SMARCA4, SMARCA5, BAZ1A, SUZ12, RBBP7). These factors may support MMR complexes during DNA repair and/or function generally in maintenance of genome stability.

We next examined how HTT KO could influence MMR interactions. An MMR-centric network was constructed with edge connections representing only Tapioca-predicted interactions (Figure 6E). This representation shows predicted interactions that are shared (Figure 6E, diamond and hexagonal nodes) and unique (Figure 6E, circular nodes). Shared interactions were represented more prominently within the MutS complex cluster (Figure 6E, green nodes). Among the most shared interactions between distinct MMR complexes (Figure 6E, hexagonal nodes), most were related to double-strand break recognition or repair (TOP2A, TOP2B, RAD50, DDX1). Outside of the known MMR interactions with MSH2 and MSH6, MSH3 had only three other interactions (RAD18, RAD50, and SMARCA5). These interactions were also shared with MSH2 and MSH6, and not with other MMR components except for a RAD18-EXO1 interaction. PMS2 interactions were only with other MMR components (Figure 6E, black edges).

To evaluate the effect of HTT KO, the change in average Tapioca score (ΔTapioca) between KO and control for a protein pair was used as a proxy for a change in their interaction. HTT KO did not appear to significantly impact core MMR-MMR interactions or cause widespread changes in interaction stability (median ΔTapioca = 0.05). However, across the MMR proteins, MLH1 interactions were notable; its 28 non-MMR interactions had a ΔTapioca score ≥ 0, suggesting many of its interactions were preferentially stabilized by HTT KO. Consistent with this observation, after filtering all MMR interactions with ΔTapioca > |0.2|, six out of the 12 HTT-dependent interactions involved MLH1, followed closely by PMS1 (4 / 12) (Supplemental Data 7). In terms of magnitude, the two predicted interactions with PMS1 (HNRNPA1 and NUPL2) were the most destabilized by HTT KO, while the most stabilized interactions were between MSH2 and TRIM33, and MLH1 and PARP2. While the significance of these changes remains to be defined, these experiments provide evidence that HTT lowering has measurable effects on the interactomes of MMR complexes.

## Discussion & Conclusions

DNA repair and handling (R/H) proteins have therapeutic promise through their roles in influencing somatic expansion and potentially disease progression (12, 14, 15, 17, 73, 74), yet the mechanisms underlying their disease modifying effects are only partially understood. Moreover, recent evidence from individuals with HD supports cell type-specific transcriptional programming differences in MMR genes (5, 6, 22), but corresponding protein patterns have not been examined. Towards these gaps in knowledge, our report established signature peptides and validated complementary workflows for the targeted mass spectrometry-based quantification of over 40 proteins involved in DNA R/H pathways. We also prioritized a subset of 11 human targets based on therapeutic interest, to demonstrate an absolute quantification assay using stable isotope dilution. The exquisite sensitivity of this assay (down to 8 amol on column) was driven by PRM-PASEF analysis on the Bruker timsTOF platform. While our assay is compatible with many instrument platforms (see discussion below), our report establishes the foundation to quantify these therapeutic candidates in especially challenging samples. For example, the assay could be deployed to establish the landscape of MMR tone in vulnerable and resilient cells from HD brains, as well as in plasma and brain-derived extracellular vesicles from CSF, which are characterized by limited sample amounts, low target levels, and wide dynamic range (75, 76). Moreover, the targeted assays developed here are broadly applicable beyond the HD models used in this study. This would accelerate mechanistic and therapeutic efforts to modulate MMR levels and activities in perturbation studies with HD models. Specifically, our developed DNA-R/H assays could be coupled with pharmacodynamic biomarker assays that validate target engagement or support proof of mechanism following ASO or RNAi knockdown (77) or targeted protein degradation/PROTAC approaches (78).

### General applications and limitations of targeted assays

In label-free workflows, we showed that the quantitative performance of both DIA and PRM-based assays on the timsTOF-based platforms was high quality, with a slight advantage for PRM-based assays of improved reproducibility and likely specificity. Our reported workflows can be performed on a variety of instrumentation. For lower throughput platforms, we generated a recommended panel of curated signature peptides that can be directly used to create instrument-specific targeted methods. However, if different sample preparation techniques are used compared to this study, not all signature peptides may be detected. Also, additional target optimization may be required when implementing the assay on platforms that we did not use in this study, e.g., triple quadrupole, tripleTOF, or Astral platforms. In this scenario, users can leverage the experiment-specific spectral libraries we assembled using FragPipe (34) (see Figure 1F and MassIVE MSV000102602) together with the Skyline assay template documents (https://panoramaweb.org/DNARH_PRM.url) to readily screen for different signature peptides as needed.

In isotope-labeled workflows, absolute quantification assays have traditionally been used for accurate measurement of sample concentrations. However, this assay could also be used for improved precision of relative (fold-change) quantification, particularly between samples with different proteome compositions and/or dynamic ranges. These issues are mitigated by using heavy isotope labeled internal reference peptides. Previous reports of MutSβ RNA and protein levels showed significant variation across diverse mouse tissues (79). Therefore, our isotope-labeled assay could be useful to provide multiplexed quantification of MMR complexes or broader DNA R/H pathways. Of note, while we developed the absolute quantification assay for 11 human proteins, 13 of the signature peptide sequences were identical between the human and mouse proteins, translating to a theoretical ability to quantify up to 7 targets in mouse samples.

While absolute quantification assays are considered the gold-standard for accurate and precise measurements, there are several assumptions that could influence quantification accuracy. First, it is possible that the total protein measurement variance contributes to the target protein’s concentration variance. It is also possible that differential proteolytic digestion also impacts protein concentration accuracy. Our assays lessened these effects by streamlining sample preparation steps, as well as selecting multiple signature peptides for each protein. However, as synthetic tryptic peptides are spiked in post-digestion, we cannot rule out the possibility that our reported protein concentrations underestimate the true values. It is theoretically possible to generate isotope-labeled recombinant protein standards, either individually or as a single protein concatemer of synthetic peptides using bacterial cell-based systems, termed QconCATs (80), or more recently, cell-free systems (81, 82). While cost effective, the production and validation of these protein standards can be challenging (83). Another approach that mitigates incomplete digestion bias involves the synthesis of “winged” peptide standards that maintain the native N- and C- trypsin cleavage sites (84). These peptides would be spiked into the samples prior to digestion and would partially compensate for differences in digestion efficiency, though this would increase the assay’s costs. For these reasons, the use of fully tryptic synthetic peptides remains the most commonly employed reagents for absolute quantification (31, 85), enabling precise determination of protein concentrations for (i) samples with high complexity and/or dynamic range, such as plasma or CSF (86–89), (ii) low abundance protein variants (90–92), or (iii) macromolecular complex stoichiometries (93, 94).

### DNA R/H protein levels in the striatum of Q140 HD mice

Our quantitative measurements of DNA R/H targets in the striatum of HD mouse models suggested minimal effects on steady-state abundance or subcellular distribution induced by mutant HTT-Q140 at the two disease stages evaluated (8 week and 40 week ages). In contrast, the nuclear versus cytoplasmic distribution of HTT and HAP40 shifted towards greater nuclear association in the HD manifest stage of the Q140 model. The accumulation of HTT fragments in various forms in the nucleus is well documented in different HD mouse models (58, 95–98). While HTT exon 1 fragments are the primary species found in nuclear inclusions and aggregates, in this study we largely measured soluble HTT, with peptide coverage that supports full-length HTT. While HTT has been shown to have a functional NLS (99) and localized to the nucleus of various mammalian cells by biochemical fractionation and immunofluorescence using multiple antibodies (100), the ability of full-length mHTT to shuttle into the nucleus is debated and may depend on specific cellular states (100, 101). While our results cannot definitively establish that full-length HTT localizes to the nuclear lumen of medium spiny neurons in mouse brain, they do suggest that a subset of mHTT relocalizes from the cytoplasm to nuclear-associated sites, such as the nuclear membrane, as the disease progresses. This is plausible given its well-established organelle and plasma membrane localizations (101, 102), as well as the ability of mHTT to disrupt the nuclear pore proteins and generally nucleocytoplasmic transport (103, 104).

### DNA R/H protein levels and interactions in human neuroblastoma cells lacking HTT

Development and application of our targeted assays in a human neuroblastoma cell line, BE(2)-C, allowed us to evaluate the impact of HTT KO on DNA R/H protein and interaction levels. We observed a general trend for downregulation for several DNA R/H proteins (Figure 5E). We cannot rule out that indirect mechanisms could contribute to global lowering of DNA R/H protein levels (105–110); nonetheless, certain targets were more strongly reduced than others. Apart from the expected decrease in HTT and HAP40, the most down-regulated protein was EXO1. The potential ability of HTT to regulate EXO1 levels, directly or indirectly, is intriguing given the recent report that wild-type HTT can interact with EXO1 and MutLα stabilizing MLH1 in the latter (111). If HTT is in a complex with these proteins, the absence of HTT may destabilize the interaction and cause degradation of EXO1. Mechanistic experiments outside of the scope of this study would be required to further delineate this relationship under HTT loss states.

We also explored consequences of HTT KO that were independent of protein expression. Thermal proximity coaggregation profiling in conjunction with Tapioca scoring, showed that the loss of HTT most strongly reshaped interactions involving MutL compared to MutS complexes (Figure 6E). Specifically, our data supports that MLH1 associations are the most stabilized due to HTT KO. A recent report from Fishwick and colleagues characterized two distinct interaction sites in the MLH1 C-terminal domain, one mediating heterodimerization with PMS2 and the other binding MIP-box motifs in EXO1, MSH3, and FAN1, which together support MutLα MMR activity. (112). Based on our TPCA experiments, HTT loss does not appear to disrupt these interactions. However, the other 28 non-MMR MLH1 interactions that we assigned have not been previously reported. Additional experiments would be required to determine whether these proteins are direct, physical interactions and/or contribute to the MMR activity of MutL.

Compared to MLH1, PMS1 interactions had more mixed stability effects due to HTT KO. For example, PMS1 showed destabilized interactions with two RNA/nuclear transport-associated proteins, HNRNPA1 and NUPL2. We found no prior evidence in the literature or in curated interaction databases connecting these proteins to PMS1, MutLα, or any other core MMR component. HNRNPA1 is an RNA-binding protein with documented affinity for G-quadruplex-forming repeat RNA structures at telomeres (113), raising the possibility of an analogous role at expanded CAG-repeat transcripts. NUPL2 (CG1) is a nucleoporin implicated in nuclear trafficking. While a specific connection of NUPL2 to DNA repair or HTT biology has not been shown, nuclear pore complexes have previously been shown to participate in genome stability (114). Moreover, as discussed above, HTT itself can localize to the nuclear membrane and has been shown to facilitate nucleocytoplasmic trafficking. We also observed a stabilized interaction between MSH2 and the E3 ubiquitin ligase TRIM33. TRIM33 has known roles in genome maintenance biology; it was recently shown to ubiquitinate and limit chromatin engagement of the transcription factor E2F4, thereby restraining E2F4-dependent recruitment of the RECQL helicase to replicating chromatin (115). Loss of TRIM33 accelerated replication under stress and impaired DNA repair (115). While this places TRIM33 in a replication stress/ubiquitin ligase functional network and not directly in an MMR network, it is plausible that HTT loss increases replication-associated stress signaling that is sensed, directly or indirectly, through MSH2, consistent with MutSα’s established role in monitoring the replication fork (116).

## Acknowledgements

We thank the CHDI Foundation for funding (A-18331), as well as contributions from NIH NIAID AI174515, NIH NIGMS GM114141, and Allen Family Philanthropies towards mass spectrometry instrumentation.

## Data Availability

The datasets produced or used in this study are available in the following public repositories:

1. Proteomics source data generated in this study have been deposited at MassIVE MSV000102602.
2. Targeted-MS source and processed data generated in this study from stable isotope dilution absolute quantification assays have been deposited in PanoramaWeb (https://panoramaweb.org/DNARH_PRM.url) and cross-listed at ProteomeXchange with the identifier PXD081801.
3. HD mouse striatum spectral library and BE(2)-C TPCA raw data used in this study: MassIVE MSV000094509, cross-listed at ProteomeXchange with the identifier PXD051339.

## Author Contributions

**Todd Greco**: Conceptualization, Formal analysis, Methodology, Software, Visualization, Validation, Writing – Original draft, Writing - Review & Editing, Project Administration, Funding acquisition **Josiah Hutton**: Conceptualization, Formal analysis, Investigation, Methodology, Resources, Visualization, Writing – Original draft. **Joshua Justice**: Investigation **Tavis Reed**: Formal analysis **Tom Vogt**: Conceptualization, Writing - Review & Editing, Funding acquisition **Brinda Prasad**: Conceptualization, Funding acquisition, Project administration, Writing – Review and Editing **Ileana Cristea**: Conceptualization, Supervision, Project administration, Funding acquisition, Writing – Review & Editing

## References

1. Bates, G. P., Dorsey, R., Gusella, J. F., Hayden, M. R., Kay, C., Leavitt, B. R., Nance, M., Ross, C. A., Scahill, R. I., Wetzel, R., Wild, E. J., and Tabrizi, S. J. (2015) Huntington disease. Nat. Rev. Dis. Primers. 10.1038/nrdp.2015.5

2. Lee, J. M., Wheeler, V. C., Chao, M. J., Vonsattel, J. P. G., Pinto, R. M., Lucente, D., Abu-Elneel, K., Ramos, E. M., Mysore, J. S., Gillis, T., MacDonald, M. E., Gusella, J. F., Harold, D., Stone, T. C., Escott-Price, V., Han, J., Vedernikov, A., Holmans, P., Jones, L., Kwak, S., Mahmoudi, M., Orth, M., Landwehrmeyer, G. B., Paulsen, J. S., Dorsey, E. R., Shoulson, I., and Myers, R. H. (2015) Identification of Genetic Factors that Modify Clinical Onset of Huntington’s Disease. Cell. 162, 516–526

3. Lee, J. M., Correia, K., Loupe, J., Kim, K. H., Barker, D., Hong, E. P., Chao, M. J., Long, J. D., Lucente, D., Vonsattel, J. P. G., Pinto, R. M., Abu Elneel, K., Ramos, E. M., Mysore, J. S., Gillis, T., Wheeler, V. C., MacDonald, M. E., Gusella, J. F., McAllister, B., Massey, T., Medway, C., Stone, T. C., Hall, L., Jones, L., Holmans, P., Kwak, S., Ehrhardt, A. G., Sampaio, C., Ciosi, M., Maxwell, A., Chatzi, A., Monckton, D. G., Orth, M., Landwehrmeyer, G. B., Paulsen, J. S., Dorsey, E. R., Shoulson, I., and Myers, R. H. (2019) CAG Repeat Not Polyglutamine Length Determines Timing of Huntington’s Disease Onset. Cell. 178, 887–900.e14

4. Kacher, R., Lejeune, F. X., Noël, S., Cazeneuve, C., Brice, A., Humbert, S., and Durr, A. (2021) Propensity for somatic expansion increases over the course of life in huntington disease. Elife. 10.7554/ELIFE.64674

5. Mätlik, K., Baffuto, M., Kus, L., Deshmukh, A. L., Davis, D. A., Paul, M. R., Carroll, T. S., Caron, M. C., Masson, J. Y., Pearson, C. E., and Heintz, N. (2024) Cell-type-specific CAG repeat expansions and toxicity of mutant Huntingtin in human striatum and cerebellum. Nat. Genet. 56, 383–394

6. Pressl, C., Mätlik, K., Kus, L., Darnell, P., Luo, J. D., Paul, M. R., Weiss, A. R., Liguore, W., Carroll, T. S., Davis, D. A., McBride, J., and Heintz, N. (2024) Selective vulnerability of layer 5a corticostriatal neurons in Huntington’s disease. Neuron. 112, 924–941.e10

7. Handsaker, R. E., Kashin, S., Reed, N. M., Tan, S., Lee, W. S., McDonald, T. M., Morris, K., Kamitaki, N., Mullally, C. D., Morakabati, N. R., Goldman, M., Lind, G., Kohli, R., Lawton, E., Hogan, M., Ichihara, K., Berretta, S., and McCarroll, S. A. (2025) Long somatic DNA-repeat expansion drives neurodegeneration in Huntington’s disease. Cell. 188, 623–639.e19

8. Farag, M., Tabrizi, S. J., and Wild, E. J. (2025) Huntington’s disease clinical trials update: October 2025. J. Huntingtons Dis. 10.1177/18796397251399751;SUBPAGE:STRING:FULL

9. Moss, D. J. H., Tabrizi, S. J., Mead, S., Lo, K., Pardiñas, A. F., Holmans, P., Jones, L., Langbehn, D., Coleman, A., Santos, R. D., Decolongon, J., Sturrock, A., Bardinet, E., Ret, C. J., Justo, D., Lehericy, S., Marelli, C., Nigaud, K., Valabrègue, R., van den Bogaard, S. J. A., Dumas, E. M., van der Grond, J., T’Hart, E. P., Jurgens, C., Witjes-Ane, M. N., Arran, N., Callaghan, J., Stopford, C., Frost, C., Jones, R., Hobbs, N., Lahiri, N., Ordidge, R., Owen, G., Pepple, T., Read, J., Say, M., Wild, E., Patel, A., Fox, N. C., Gibbard, C., Malone, I., Crawford, H., Whitehead, D., Keenan, S., Cash, D. M., Berna, C., Bechtel, N., Bohlen, S., Man, A. H., Kraus, P., Axelson, E., Wang, C., Acharya, T., Lee, S., Monaco, W., Campbell, C., Queller, S., Whitlock, K., Campbell, C., Campbell, M., Frajman, E., Milchman, C., O’Regan, A., Labuschagne, I., Stout, J., Landwehrmeyer, B., Craufurd, D., Scahill, R., Hicks, S., Kennard, C., Johnson, H., Tobin, A., Rosas, H. D., Reilmann, R., Borowsky, B., Pourchot, C., Andrews, S. C., Bachoud-Lévi, A. C., Bentivoglio, A. R., Biunno, I., Bonelli, R., Burgunder, J. M., Dunnett, S., Ferreira, J., Handley, O., Heiberg, A., Illmann, T., Landwehrmeyer, G. B., Levey, J., Ramos-Arroyo, M. A., Nielsen, J., Koivisto, S. P., Päivärinta, M., Roos, R. A. C., Sebastián, A. R., Tabrizi, S., Vandenberghe, W., Verellen-Dumoulin, C., Uhrova, T., Wahlström, J., Zaremba, J., Baake, V., Barth, K., Garde, M. B., Betz, S., Bos, R., Callaghan, J., Come, A., Guedes, L. C., Ecker, D., Finisterra, A. M., Fullam, R., Gilling, M., Gustafsson, L., Handley, O. J., Hvalstedt, C., Held, C., Koppers, K., Lamanna, C., Laurà, M., Descals, A. M., Martinez-Horta, S., Mestre, T., Minster, S., Monza, D., Mütze, L., Oehmen, M., Orth, M., Padieu, H., Paterski, L., Peppa, N., Koivisto, S. P., Di Renzo, M., Rialland, A., Røren, N., Šašinková, P., Timewell, E., Townhill, J., Cubillo, P. T., da Silva, W. V., van Walsem, M. R., Whalstedt, C., Witjes-Ané, M. N., Witkowski, G., Wright, A., Zielonka, D., Zielonka, E., Zinzi, P., Bonelli, R. M., Lilek, S., Hecht, K., Herranhof, B., Holl, A., Kapfhammer, H. P., Koppitz, M., Magnet, M., Müller, N., Otti, D., Painold, A., Reisinger, K., Scheibl, M., Schöggl, H., Ullah, J., Braunwarth, E. M., Brugger, F., Buratti, L., Hametner, E. M., Hepperger, C., Holas, C., Hotter, A., Hussl, A., Müller, C., Poewe, W., Seppi, K., Sprenger, F., Wenning, G., Boogaerts, A., Calmeyn, G., Delvaux, I., Liessens, D., Somers, N., Dupuit, M., Minet, C., van Paemel, D., Ribaï, P., Verellen-Dumoulin, C., Boogaerts, A., Vandenberghe, W., van Reijen, D., Klempír, J., Majerová, V., Roth, J., Stárková, I., Hjermind, L. E., Jacobsen, O., Nielsen, J. E., Larsen, I. U., Vinther-Jensen, T., Hiivola, H., Hyppönen, H., Martikainen, K., Tuuha, K., Allain, P., Bonneau, D., Bost, M., Gohier, B., Guérid, M. A., Olivier, A., Prundean, A., Scherer-Gagou, C., Verny, C., Babiloni, B., Debruxelles, S., Duché, C., Goizet, C., Jameau, L., Lafoucrière, D., Spampinato, U., Barthélémy, R., De Bruycker, C., Carette, M. C. A. S., Defebvre, E. D. L., Delliaux, M., Delval, A., Destee, A., Dujardin, K., Lemaire, M. H., Manouvrier, S., Peter, M., Plomhouse, L., Sablonnière, B., Simonin, C., Thibault-Tanchou, S., Vuillaume, I., Bellonet, M., Berrissoul, H., Blin, S., Courtin, F., Duru, C., Fasquel, V., Godefroy, O., Krystkowiak, P., Mantaux, B., Roussel, M., Wannepain, S., Azulay, J. P., Delfini, M., Eusebio, A., Fluchere, F., Mundler, L., Anheim, M., Julié, C., Boukbiza, O. L., Longato, N., Rudolf, G., Tranchant, C., Zimmermann, M. A., Kosinski, C. M., Milkereit, E., Probst, D., Reetz, K., Sass, C., Schiefer, J., Schlangen, C., Werner, C. J., Gelderblom, H., Priller, J., Prüß, H., Spruth, E. J., Ellrichmann, G., Herrmann, L., Hoffmann, R., Kaminski, B., Kotz, P., Prehn, C., Saft, C., Lange, H., Maiwald, R., Löhle, M., Maass, A., Schmidt, S., Bosredon, C., Storch, A., Wolz, A., Wolz, M., Capetian, P., Lambeck, J., Zucker, B., Boelmans, K., Ganos, C., Heinicke, W., Hidding, U., Lewerenz, J., Münchau, A., Orth, M., Schmalfeld, J., Stubbe, L., Zittel, S., Diercks, G., Dressler, D., Gorzolla, H., Schrader, C., Tacik, P., Ribbat, M., Longinus, B., Bürk, K., Möller, J. C., Rissling, I., Mühlau, M., Peinemann, A., Städtler, M., Weindl, A., Winkelmann, J., Ziegler, C., Bechtel, N., Beckmann, H., Bohlen, S., Hölzner, E., Lange, H., Reilmann, R., Rohm, S., Rumpf, S., Schepers, S., Weber, N., Dose, M., Leythäuser, G., Marquard, R., Raab, T., Wiedemann, A., Barth, K., Buck, A., Connemann, J., Ecker, D., Geitner, C., Held, C., Kesse, A., Landwehrmeyer, B., Lang, C., Lewerenz, J., Lezius, F., Nepper, S., Niess, A., Orth, M., Schneider, A., Schwenk, D., Süßmuth, S., Trautmann, S., Weydt, P., Cormio, C., Sciruicchio, V., Serpino, C., de Tommaso, M., Capellari, S., Cortelli, P., Galassi, R., Rizzo, G., Poda, R., Scaglione, C., Bertini, E., Ghelli, E., Ginestroni, A., Massaro, F., Mechi, C., Paganini, M., Piacentini, S., Pradella, S., Romoli, A. M., Sorbi, S., Abbruzzese, G., di Poggio, M. B., Ferrandes, G., Mandich, P., Marchese, R., Albanese, A., Di Bella, D., Castaldo, A., Di Donato, S., Gellera, C., Genitrini, S., Mariotti, C., Monza, D., Nanetti, L., Paridi, D., Soliveri, P., Tomasello, C., De Michele, G., Di Maio, L., Massarelli, M., Peluso, S., Roca, A., Russo, C. V., Salvatore, E., Sorrentino, P., Amico, E., Favellato, M., Griguoli, A., Mazzante, I., Petrollini, M., Squitieri, F., D’Alessio, B., Esposito, C., Bentivoglio, R., Frontali, M., Guidubaldi, A., Ialongo, T., Jacopini, G., Piano, C., Romano, S., Soleti, F., Spadaro, M., Zinzi, P., van Hout, M. S. E., Verhoeven, M. E., van Vugt, J. P. P., de Weert, A. M., Bolwijn, J. J. W., Dekker, M., Kremer, B., Leenders, K. L., van Oostrom, J. C. H., van den Bogaard, S. J. A., Bos, R., Dumas, E. M., ’t Hart, E. P., Roos, R. A. C., Kremer, B., Verstappen, C. C. P., Aaserud, O., Jan Frich, C., Heiberg, A., van Walsem, M. R., Wehus, R., Bjørgo, K., Fannemel, M., Gørvell, P. F., Lorentzen, E., Koivisto, S. P., Retterstøl, L., Stokke, B., Bjørnevoll, I., Sando, S. B., Dziadkiewicz, A., Nowak, M., Robowski, P., Sitek, E., Slawek, J., Soltan, W., Szinwelski, M., Blaszcyk, M., Boczarska-Jedynak, M., Ciach-Wysocka, E., Gorzkowska, A., Jasinska-Myga, B., Klodowska-Duda, G., Opala, G., Stompel, D., Banaszkiewicz, K., Bocwinska, D., Bojakowska-Jaremek, K., Dec, M., Krawczyk, M., Rudzinska, M., Szczygiel, E., Szczudlik, A., Wasielewska, A., Wójcik, M., Bryl, A., Ciesielska, A., Klimberg, A., Marcinkowski, J., Samara, H., Sempolowicz, J., Zielonka, D., Gogol, A., Janik, P., Kwiecinski, H., Jamrozik, Z., Antczak, J., Jachinska, K., Krysa, W., Rakowicz, M., Richter, P., Rola, R., Ryglewicz, D., Sienkiewicz-Jarosz, H., Stepniak, I., Sulek, A., Witkowski, G., Zaremba, J., Zdzienicka, E., Zieora-Jakutowicz, K., Ferreira, J. J., Coelho, M., Guedes, L. C., Mendes, T., Mestre, T., Valadas, A., Andrade, C., Gago, M., Garrett, C., Guerra, M. R., Herrera, C. D., Garcia, P. M., Barbera, M. A., Guia, D. B., Hernanz, L. C., Catena, J. L., Ferrer, P. Q., Sebastián, A. R., Carruesco, G. T., Bas, J., Busquets, N., Calopa, M., Robert, M. F., Viladrich, C. M., Idiago, J. M. R., Riballo, A. V., Cubo, E., Polo, C. G., Mariscal, N., Rivadeneyra, P. J., Barrero, F., Morales, B., Fenollar, M., García, R. G. R., Ortega, P., Villanueva, C., Alegre, J., Bascuñana, M., Caldentey, J. G., Ventura, M. F., Ribas, G. G., de Yébenes, J. G., Moreno, J. L. L. S., Cubillo, P. T., Alegre, J., Frech, F. A., de Yébenes, J. G., Ruíz, P. J. G., Martínez-Descals, A., Guerrero, R., Artiga, M. J. S., Sánchez, V., Perea, M. F. N., Fortuna, L., Manzanares, S., Reinante, G., Torres, M. M. A., Moreau, L. V., González González, S., Guisasola, L. M., Salvador, C., Martín, E. S. S., Ramirez, I. L., Gorospe, A., Lopera, M. R., Arques, P. N., Rodríguez, M. J. T., Pastor, B. V., Gaston, I., Martinez-Jaurrieta, M. D., Ramos-Arroyo, M. A., Moreno, J. M. G., Lucena, C. M., Damas, F., Cortegana, H. E. P., Peña, J. C., Redondo, L., Carrillo, F., Teresa Cáceres, M., Mir, P., Suarez, M. J. L., Vargas-González, L., Bosca, M. E., Brugada, F. C., Burguera, J. A., Campos, A., Vilaplana, G. C. P., Berglund, P., Constantinescu, R., Fredlund, G., Høsterey-Ugander, U., Linnsand, P., Neleborn-Lingefjärd, L., Wahlström, J., Wentzel, M., Loutfi, G., Olofsson, C., Stattin, E. L., Westman, L., Wikström, B., Burgunder, J. M., Stebler, Y., Kaelin, A., Romero, I., Schüpbach, M., Weber Zaugg, S., Hauer, M., Gonzenbach, R., Jung, H. H., Mihaylova, V., Petersen, J., Jack, R., Matheson, K., Miedzybrodzka, Z., Rae, D., Simpson, S. A., Summers, F., Ure, A., Vaughan, V., Akhtar, S., Crooks, J., Curtis, A., de Souza, J., Piedad, J., Rickards, H., Wright, J., Coulthard, E., Gethin, L., Hayward, B., Sieradzan, K., Wright, A., Armstrong, M., Barker, R. A., O’Keefe, D., Di Pietro, A., Fisher, K., Goodman, A., Hill, S., Kershaw, A., Mason, S., Paterson, N., Raymond, L., Swain, R., Guzman, N. V., Busse, M., Butcher, C., Callaghan, J., Dunnett, S., Clenaghan, C., Fullam, R., Handley, O., Hunt, S., Jones, L., Jones, U., Khalil, H., Minster, S., Owen, M., Price, K., Rosser, A., Townhill, J., Edwards, M., Ho, C., Hughes, T., McGill, M., Pearson, P., Porteous, M., Smith, P., Brockie, P., Foster, J., Johns, N., McKenzie, S., Rothery, J., Thomas, G., Yates, S., Burrows, L., Chu, C., Fletcher, A., Gallantrae, D., Hamer, S., Harding, A., Klöppel, S., Kraus, A., Laver, F., Lewis, M., Longthorpe, M., Markova, I., Raman, A., Robertson, N., Silva, M., Thomson, A., Wild, S., Yardumian, P., Chu, C., Evans, C., Gallentrae, D., Hamer, S., Kraus, A., Markova, I., Raman, A., Chu, C., Hamer, S., Hobson, E., Jamieson, S., Kraus, A., Markova, I., Raman, A., Musgrave, H., Rowett, L., Toscano, J., Wild, S., Yardumian, P., Bourne, C., Clapton, J., Clayton, C., Dipple, H., Freire-Patino, D., Grant, J., Gross, D., Hallam, C., Middleton, J., Murch, A., Thompson, C., Alusi, S., Davies, R., Foy, K., Gerrans, E., Pate, L., Andrews, T., Dougherty, A., Golding, C., Kavalier, F., Laing, H., Lashwood, A., Robertson, D., Ruddy, D., Santhouse, A., Whaite, A., Andrews, T., Bruno, S., Doherty, K., Golding, C., Haider, S., Hensman, D., Lahiri, N., Lewis, M., Novak, M., Patel, A., Robertson, N., Rosser, E., Tabrizi, S., Taylor, R., Warner, T., Wild, E., Arran, N., Bek, J., Callaghan, J., Craufurd, D., Fullam, R., Hare, M., Howard, L., Huson, S., Johnson, L., Jones, M., Murphy, H., Oughton, E., Partington-Jones, L., Rogers, D., Sollom, A., Snowden, J., Stopford, C., Thompson, J., Trender-Gerhard, I., Verstraelen, N., Westmoreland, L., Armstrong, R., Dixon, K., Nemeth, A. H., Siuda, G., Valentine, R., Harrison, D., Hughes, M., Parkinson, A., Soltysiak, B., Bandmann, O., Bradbury, A., Gill, P., Fairtlough, H., Fillingham, K., Foustanos, I., Kazoka, M., O’Donovan, K., Peppa, N., Taylor, C., Tidswell, K., Quarrell, O., Burgunder, J. M., Lau, P. N., Pica, E., and Tan, L. (2017) Identification of genetic variants associated with Huntington’s disease progression: a genome-wide association study. Lancet Neurol. 16, 701–711

10. Myers, R. H., Dorsey, E. R., Paulsen, J. S., Landwehrmeyer, G. B., Orth, M., Sampaio, C., Kwak, S., Holmans, P., Jones, L., Massey, T. H., Wills, C., Monckton, D. G., Loay, H., Lomeikaite, V., Ciosi, M., Gatseva, A., Gusella, J. F., MacDonald, M. E., Wheeler, V. C., Gillis, T., Ruliera, J., Elezi, E., Siciliano, J., Mysore, J. S., Giordano, J. V., Mouro Pinto, R., Seong, I. S., Lucente, D., Long, J. D., Choi, D. E., Kim, K. H., Lee, Y., Jang, J. H., Lee, S., Shin, J. W., Correia, K., McLean, Z. L., and Lee, J. M. (2025) Genetic modifiers of somatic expansion and clinical phenotypes in Huntington’s disease highlight shared and tissue-specific effects. Nat. Genet. 57, 1426–1436

11. Dragileva, E., Hendricks, A., Teed, A., Gillis, T., Lopez, E. T., Friedberg, E. C., Kucherlapati, R., Edelmann, W., Lunetta, K. L., MacDonald, M. E., and Wheeler, V. C. (2009) Intergenerational and striatal CAG repeat instability in Huntington’s disease knock-in mice involve different DNA repair genes. Neurobiol. Dis. 33, 37–47

12. Tomé, S., Manley, K., Simard, J. P., Clark, G. W., Slean, M. M., Swami, M., Shelbourne, P. F., Tillier, E. R. M., Monckton, D. G., Messer, A., and Pearson, C. E. (2013) MSH3 Polymorphisms and Protein Levels Affect CAG Repeat Instability in Huntington’s Disease Mice. PLoS Genet. 9, e1003280

13. Keogh, N., Chan, K. Y., Li, G. M., and Lahue, R. S. (2017) MutSβ abundance and Msh3 ATP hydrolysis activity are important drivers of CTG•CAG repeat expansions. Nucleic Acids Res. 45, 10068–10078

14. Driscoll, R., Hampton, L., Abraham, N. A., Larigan, J. D., Joseph, N. F., Hernandez-Vega, J. C., Geisler, S., Yang, F. C., Deninger, M., Tran, D. T., Khatri, N., Godinho, B. M. D. C., Kinberger, G. A., Montagna, D. R., Hirst, W. D., Guardado, C. L., Glajch, K. E., Arnold, H. M., Gallant-Behm, C. L., and Weihofen, A. (2024) Dose-dependent reduction of somatic expansions but not Htt aggregates by di-valent siRNA-mediated silencing of MSH3 in HdhQ111 mice. Scientific Reports 2024 14:1. 14, 2061-

15. Wang, N., Zhang, S., Langfelder, P., Ramanathan, L., Gao, F., Plascencia, M., Vaca, R., Gu, X., Deng, L., Dionisio, L. E., Vu, H., Maciejewski, E., Ernst, J., Prasad, B. C., Vogt, T. F., Horvath, S., Aaronson, J. S., Rosinski, J., and Yang, X. W. (2025) Distinct mismatch-repair complex genes set neuronal CAG-repeat expansion rate to drive selective pathogenesis in HD mice. Cell. 188, 1524–1544.e22

16. Aldous, S. G., Smith, E. J., Landles, C., Osborne, G. F., Cañibano-Pico, M., Nita, I. M., Phillips, J., Zhang, Y., Jin, B., Hirst, M. B., Benn, C. L., Bond, B. C., Edelmann, W., Greene, J. R., and Bates, G. P. (2024) A CAG repeat threshold for therapeutics targeting somatic instability in Huntington’s disease. Brain. 147, 1784–1798

17. Pinto, R. M., Dragileva, E., Kirby, A., Lloret, A., Lopez, E., St. Claire, J., Panigrahi, G. B., Hou, C., Holloway, K., Gillis, T., Guide, J. R., Cohen, P. E., Li, G. M., Pearson, C. E., Daly, M. J., and Wheeler, V. C. (2013) Mismatch repair genes Mlh1 and Mlh3 modify CAG instability in Huntington’s disease mice: Genome-wide and candidate approaches. PLoS Genet. 10.1371/journal.pgen.1003930

18. Loupe, J. M., Pinto, R. M., Kim, K. H., Gillis, T., Mysore, J. S., Andrew, M. A., Kovalenko, M., Murtha, R., Seong, I., Gusella, J. F., Kwak, S., Howland, D., Lee, R., Lee, J. M., Wheeler, V. C., and MacDonald, M. E. (2020) Promotion of somatic CAG repeat expansion by Fan1 knock-out in Huntington’s disease knock-in mice is blocked by Mlh1 knock-out. Hum. Mol. Genet. 29, 3044–3053

19. Goold, R., Flower, M., Moss, D. H., Medway, C., Wood-Kaczmar, A., Andre, R., Farshim, P., Bates, G. P., Holmans, P., Jones, L., and Tabrizi, S. J. (2019) FAN1 modifies Huntington’s disease progression by stabilizing the expanded HTT CAG repeat. Hum. Mol. Genet. 28, 650–661

20. Goold, R., Hamilton, J., Menneteau, T., Flower, M., Bunting, E. L., Aldous, S. G., Porro, A., Vicente, J. R., Allen, N. D., Wilkinson, H., Bates, G. P., Sartori, A. A., Thalassinos, K., Balmus, G., and Tabrizi, S. J. (2021) FAN1 controls mismatch repair complex assembly via MLH1 retention to stabilize CAG repeat expansion in Huntington’s disease. Cell Rep. 10.1016/j.celrep.2021.109649

21. Bunting, E. L., Panhale, A., McColgan, P., Koi, M., Brundin, P., and Carethers, J. M. (2026) Targeting DNA mismatch repair in Huntington’s disease. Trends Neurosci. 10.1016/j.tins.2026.06.004

22. Baffuto, M., Mätlik, K., Ilyashov, I., Chetia, H., Siantoputri, M. E., Sipos, E., Maeda, Y., Darnell, P., Didkovsky, N., Kus, L., Carroll, T. S., Barrows, D., Pressl, C., and Heintz, N. (2025) Epigenetic mechanisms governing cell type specific somatic expansion and toxicity in Huntington’s disease. bioRxiv. 10.1101/2025.05.21.653721

23. Harenza, J. L., DIamond, M. A., Adams, R. N., Song, M. M., Davidson, H. L., Hart, L. S., Dent, M. H., Fortina, P., Reynolds, C. P., and Maris, J. M. (2017) Transcriptomic profiling of 39 commonly-used neuroblastoma cell lines. Sci. Data. 10.1038/SDATA.2017.33,

24. Guzman, U. H., Martinez-Val, A., Ye, Z., Damoc, E., Arrey, T. N., Pashkova, A., Renuse, S., Denisov, E., Petzoldt, J., Peterson, A. C., Harking, F., Østergaard, O., Rydbirk, R., Aznar, S., Stewart, H., Xuan, Y., Hermanson, D., Horning, S., Hock, C., Makarov, A., Zabrouskov, V., and Olsen, J. V. (2024) Ultra-fast label-free quantification and comprehensive proteome coverage with narrow-window data-independent acquisition. Nat. Biotechnol. 10.1038/S41587-023-02099-7,

25. Pouladi, M. A., Morton, A. J., and Hayden, M. R. (2013) Choosing an animal model for the study of Huntington’s disease. Nat. Rev. Neurosci. 14, 708–721

26. Stocksdale, J. T., Leventhal, M. J., Lam, S., Xu, Y. X., Wang, Y. O., Wang, K. Q., Thomas, R., Faghihmonzavi, Z., Raghav, Y., Smith, C., Wu, J., Miramontes, R., Sarda, K., Johnston, H., Shin, M. G., Huang, T., Foster, M., Barch, M., Amirani, N., Paiz, C., Easter, L., Duderstadt, E., Vaibhav, V., Sundararaman, N., Felsenfeld, D. P., Vogt, T. F., Van Eyk, J., Finkbeiner, S., Kaye, J. A., Fraenkel, E., and Thompson, L. M. (2025) Intersecting impact of CAG repeat and huntingtin knockout in stem cell-derived cortical neurons. Neurobiol. Dis. 10.1016/j.nbd.2025.106914

27. Zhu, W., Smith, J. W., and Huang, C. M. (2010) Mass spectrometry-based label-free quantitative proteomics. J. Biomed. Biotechnol. 10.1155/2010/840518

28. Peterson, A. C., Russell, J. D., Bailey, D. J., Westphall, M. S., and Coon, J. J. (2012) Parallel reaction monitoring for high resolution and high mass accuracy quantitative, targeted proteomics. Molecular and Cellular Proteomics. 11, 1475–1488

29. Kennedy, M. A., Tyl, M. D., Betsinger, C. N., Federspiel, J. D., Sheng, X., Arbuckle, J. H., Kristie, T. M., and Cristea, I. M. (2022) A TRUSTED targeted mass spectrometry assay for pan-herpesvirus protein detection. Cell Rep. 10.1016/j.celrep.2022.110810

30. Wiśniewski, J. R., Hein, M. Y., Cox, J., and Mann, M. (2014) A “proteomic ruler” for protein copy number and concentration estimation without spike-in standards. Molecular and Cellular Proteomics. 13, 3497–3506

31. Gerber, S. A., Rush, J., Stemman, O., Kirschner, M. W., and Gygi, S. P. (2003) Absolute quantification of proteins and phosphoproteins from cell lysates by tandem MS. Proc. Natl. Acad. Sci. U. S. A. 100, 6940–6945

32. Kettenbach, A. N., Rush, J., and Gerber, S. A. (2011) Absolute quantification of protein and post-translational modification abundance with stable isotope-labeled synthetic peptides. Nat. Protoc. 6, 175–186

33. Li, K., Teo, G. C., Yang, K. L., Yu, F., and Nesvizhskii, A. I. (2025) diaTracer enables spectrum-centric analysis of diaPASEF proteomics data. Nat. Commun. 16, 95

34. Yu, F., Deng, Y., and Nesvizhskii, A. I. (2025) MSFragger-DDA+ enhances peptide identification sensitivity with full isolation window search. Nat. Commun. 10.1038/S41467-025-58728-Z

35. Demichev, V., Messner, C. B., Vernardis, S. I., Lilley, K. S., and Ralser, M. (2019) DIA-NN: neural networks and interference correction enable deep proteome coverage in high throughput. Nature Methods 2019 17:1. 17, 41–44

36. MacLean, B., Tomazela, D. M., Shulman, N., Chambers, M., Finney, G. L., Frewen, B., Kern, R., Tabb, D. L., Liebler, D. C., and MacCoss, M. J. (2010) Skyline: an open source document editor for creating and analyzing targeted proteomics experiments. Bioinformatics. 26, 966–968

37. Zhu, Y., Orre, L. M., Zhou Tran, Y., Mermelekas, G., Johansson, H. J., Malyutina, A., Anders, S., and Lehtiö, J. (2020) DEqMS: a method for accurate variance estimation in differential protein expression analysis. Molecular & Cellular Proteomics. 10.1074/mcp.tir119.001646

38. Justice, J. L., Kennedy, M. A., Hutton, J. E., Liu, D., Song, B., Phelan, B., and Cristea, I. M. (2021) Systematic profiling of protein complex dynamics reveals DNA-PK phosphorylation of IFI16 en route to herpesvirus immunity. Sci. Adv. 7, 6680–6698

39. Zeng, W. F., Zhou, X. X., Willems, S., Ammar, C., Wahle, M., Bludau, I., Voytik, E., Strauss, M. T., and Mann, M. (2022) AlphaPeptDeep: a modular deep learning framework to predict peptide properties for proteomics. Nat. Commun. 10.1038/S41467-022-34904-3

40. Skowronek, P., Thielert, M., Voytik, E., Tanzer, M. C., Hansen, F. M., Willems, S., Karayel, O., Brunner, A. D., Meier, F., and Mann, M. (2022) Rapid and In-Depth Coverage of the (Phospho-)Proteome With Deep Libraries and Optimal Window Design for dia-PASEF. Mol. Cell. Proteomics. 10.1016/J.MCPRO.2022.100279

41. Justice, J. L., Greco, T. M., Hutton, J. E., Reed, T. J., Mair, M. L., Botas, J., and Cristea, I. M. (2024) Multi-epitope immunocapture of huntingtin reveals striatum-selective molecular signatures. bioRxiv. 10.1101/2024.09.07.611843

42. Reed, T. J., Tyl, M. D., Tadych, A., Troyanskaya, O. G., and Cristea, I. M. (2024) Tapioca: a platform for predicting de novo protein-protein interactions in dynamic contexts. Nat. Methods. 21, 488–500

43. Lee, J. M., Huang, Y., Orth, M., Gillis, T., Siciliano, J., Hong, E., Mysore, J. S., Lucente, D., Wheeler, V. C., Seong, I. S., McLean, Z. L., Mills, J. A., McAllister, B., Lobanov, S. V., Massey, T. H., Ciosi, M., Landwehrmeyer, G. B., Paulsen, J. S., Dorsey, E. R., Shoulson, I., Sampaio, C., Monckton, D. G., Kwak, S., Holmans, P., Jones, L., MacDonald, M. E., Long, J. D., and Gusella, J. F. (2022) Genetic modifiers of Huntington disease differentially influence motor and cognitive domains. Am. J. Hum. Genet. 109, 885–899

44. Namuli, K. L., Slike, A. N., Hollebeke, M. A., and Wright, G. E. B. (2025) Genomic characterization of Huntington’s disease genetic modifiers informs drug target tractability. Brain Commun. 10.1093/BRAINCOMMS/FCAE418

45. Lobanov, S. V., McAllister, B., McDade-Kumar, M., Landwehrmeyer, G. B., Orth, M., Rosser, A. E., Paulsen, J. S., Lee, J. M., MacDonald, M. E., Gusella, J. F., Long, J. D., Ryten, M., Williams, N. M., Holmans, P., Massey, T. H., and Jones, L. (2022) Huntington’s disease age at motor onset is modified by the tandem hexamer repeat in TCERG1. NPJ Genom. Med. 10.1038/S41525-022-00317-W

46. Kim, K. H., Hong, E. P., Shin, J. W., Chao, M. J., Loupe, J., Gillis, T., Mysore, J. S., Holmans, P., Jones, L., Orth, M., Monckton, D. G., Long, J. D., Kwak, S., Lee, R., Gusella, J. F., MacDonald, M. E., and Lee, J. M. (2020) Genetic and Functional Analyses Point to FAN1 as the Source of Multiple Huntington Disease Modifier Effects. Am. J. Hum. Genet. 107, 96–110

47. Vidova, V., and Spacil, Z. (2017) A review on mass spectrometry-based quantitative proteomics: Targeted and data independent acquisition. Anal. Chim. Acta. 964, 7–23

48. Picotti, P., and Aebersold, R. (2012) Selected reaction monitoring-based proteomics: workflows, potential, pitfalls and future directions. Nat. Methods. 9, 555–566

49. Zhu, H., Ficarro, S. B., Alexander, W. M., Fleming, L. E., Adelmant, G., Zhang, T., Willetts, M., Decker, J., Brehmer, S., Krause, M., East, M. P., Gray, N. S., Johnson, G. L., Kruppa, G., and Marto, J. A. (2021) PRM-LIVE with Trapped Ion Mobility Spectrometry and Its Application in Selectivity Profiling of Kinase Inhibitors. Anal. Chem. 93, 13791–13799

50. Menalled, L. B., Sison, J. D., Dragatsis, I., Zeitlin, S., and Chesselet, M. F. (2003) Time course of early motor and neuropathological anomalies in a knock-in mouse model of Huntington’s disease with 140 CAG repeats. Journal of Comparative Neurology. 465, 11–26

51. Zheng, S., Ghitani, N., Blackburn, J. S., Liu, J. P., and Zeitlin, S. O. (2012) A series of N-terminal epitope tagged Hdh knock-in alleles expressing normal and mutant huntingtin: Their application to understanding the effect of increasing the length of normal huntingtins polyglutamine stretch on CAG140 mouse model pathogenesis. Mol. Brain. 10.1186/1756-6606-5-28

52. Guo, Q., Huang, B., Cheng, J., Seefelder, M., Engler, T., Pfeifer, G., Oeckl, P., Otto, M., Moser, F., Maurer, M., Pautsch, A., Baumeister, W., Fernández-Busnadiego, R., and Kochanek, S. (2018) The cryo-electron microscopy structure of huntingtin. Nature. 555, 117–120

53. Harding, R. J., Deme, J. C., Hevler, J. F., Tamara, S., Lemak, A., Cantle, J. P., Szewczyk, M. M., Begeja, N., Goss, S., Zuo, X., Loppnau, P., Seitova, A., Hutchinson, A., Fan, L., Truant, R., Schapira, M., Carroll, J. B., Heck, A. J. R., Lea, S. M., and Arrowsmith, C. H. (2021) Huntingtin structure is orchestrated by HAP40 and shows a polyglutamine expansion-specific interaction with exon 1. *Commun*. Biol. 10.1038/S42003-021-02895-4

54. Justice, J. L., Greco, T. M., Hutton, J. E., Reed, T. J., Mair, M. L., Botas, J., and Cristea, I. M. (2025) Multi-epitope immunocapture of huntingtin reveals striatum-selective molecular signatures. Mol. Syst. Biol. 21, 492–522

55. Langfelder, P., Cantle, J. P., Chatzopoulou, D., Wang, N., Gao, F., Al-Ramahi, I., Lu, X. H., Ramos, E. M., El-Zein, K., Zhao, Y., Deverasetty, S., Tebbe, A., Schaab, C., Lavery, D. J., Howland, D., Kwak, S., Botas, J., Aaronson, J. S., Rosinski, J., Coppola, G., Horvath, S., and Yang, X. W. (2016) Integrated genomics and proteomics define huntingtin CAG length-dependent networks in mice. Nat. Neurosci. 19, 623–633

56. Scherzinger, E., Lurz, R., Turmaine, M., Mangiarini, L., Hollenbach, B., Hasenbank, R., Bates, G. P., Davies, S. W., Lehrach, H., and Wanker, E. E. (1997) Huntingtin-encoded polyglutamine expansions form amyloid-like protein aggregates in vitro and in vivo. Cell. 90, 549–558

57. Davies, S. W., Turmaine, M., Cozens, B. A., DiFiglia, M., Sharp, A. H., Ross, C. A., Scherzinger, E., Wanker, E. E., Mangiarini, L., and Bates, G. P. (1997) Formation of neuronal intranuclear inclusions underlies the neurological dysfunction in mice transgenic for the HD mutation. Cell. 90, 537–548

58. DiFiglia, M., Sapp, E., Chase, K. O., Davies, S. W., Bates, G. P., Vonsattel, J. P., and Aronin, N. (1997) Aggregation of huntingtin in neuronal intranuclear inclusions and dystrophic neurites in brain. Science (1979). 277, 1990–1993

59. Crook, Z. R., and Housman, D. (2011) Huntington’s Disease: Can Mice Lead the Way to Treatment? Neuron. 69, 423

60. Nuzzo, M. T., Fiocchetti, M., Totta, P., Melone, M. A. B., Cardinale, A., Fusco, F. R., Gustincich, S., Persichetti, F., Ascenzi, P., and Marino, M. (2016) Huntingtin polyQ Mutation Impairs the 17β-Estradiol/Neuroglobin Pathway Devoted to Neuron Survival. Molecular Neurobiology 2016 54:8. 54, 6634–6646

61. Bensalel, J., Xu, H., Lu, M. L., Capobianco, E., and Wei, J. (2021) RNA-seq analysis reveals significant transcriptome changes in huntingtin-null human neuroblastoma cells. BMC Med. Genomics. 10.1186/S12920-021-01022-W

62. Pino, L. K., Searle, B. C., Yang, H. Y., Hoofnagle, A. N., Noble, W. S., and MacCoss, M. J. (2020) Matrix-Matched Calibration Curves for Assessing Analytical Figures of Merit in Quantitative Proteomics. J. Proteome Res. 19, 1147–1153

63. Carr, S. A., Abbatiello, S. E., Ackermann, B. L., Borchers, C., Domon, B., Deutsch, E. W., Grant, R. P., Hoofnagle, A. N., Hüttenhain, R., Koomen, J. M., Liebler, D. C., Liu, T., MacLean, B., Mani, D. R., Mansfield, E., Neubert, H., Paulovich, A. G., Reiter, L., Vitek, O., Aebersold, R., Anderson, L., Bethem, R., Blonder, J., Boja, E., Botelho, J., Boyne, M., Bradshaw, R. A., Burlingame, A. L., Chan, D., Keshishian, H., Kuhn, E., Kinsinger, C., Lee, J. S. H., Lee, S. W., Moritz, R., Oses-Prieto, J., Rifai, N., Ritchie, J., Rodriguez, H., Srinivas, P. R., Townsend, R. R., Van Eyk, J., Whiteley, G., Wiita, A., and Weintraub, S. (2014) Targeted peptide measurements in biology and medicine: Best practices for mass spectrometry-based assay development using a fit-for-purpose approach. Molecular and Cellular Proteomics. 13, 907–917

64. Beck, M., Schmidt, A., Malmstroem, J., Claassen, M., Ori, A., Szymborska, A., Herzog, F., Rinner, O., Ellenberg, J., and Aebersold, R. (2011) The quantitative proteome of a human cell line. Mol. Syst. Biol. 10.1038/MSB.2011.82,

65. Bekker-Jensen, D. B., Kelstrup, C. D., Batth, T. S., Larsen, S. C., Haldrup, C., Bramsen, J. B., Sørensen, K. D., Høyer, S., Ørntoft, T. F., Andersen, C. L., Nielsen, M. L., and Olsen, J. V. (2017) An Optimized Shotgun Strategy for the Rapid Generation of Comprehensive Human Proteomes. Cell Syst. 4, 587–599.e4

66. Macdonald, D., Tessari, M. A., Boogaard, I., Smith, M., Pulli, K., Szynol, A., Albertus, F., Lamers, M. B. A. C., Dijkstra, S., Kordt, D., Reindl, W., Herrmann, F., McAllister, G., Fischer, D. F., and Munoz-Sanjuan, I. (2014) Quantification assays for total and polyglutamine-expanded huntingtin proteins. PLoS One. 10.1371/JOURNAL.PONE.0096854,

67. Zhang, H., Das, S., Li, Q. Z., Dragatsis, I., Repa, J., Zeitlin, S., Hajnóczky, G., and Bezprozvanny, I. (2008) Elucidating a normal function of huntingtin by functional and microarray analysis of huntingtin-null mouse embryonic fibroblasts. BMC Neurosci. 9, 38

68. Tan, C. S. H., Go, K. D., Bisteau, X., Dai, L., Yong, C. H., Prabhu, N., Ozturk, M. B., Lim, Y. T., Sreekumar, L., Lengqvist, J., Tergaonkar, V., Kaldis, P., Sobota, R. M., and Nordlund, P. (2018) Thermal proximity coaggregation for system-wide profiling of protein complex dynamics in cells. Science (1979). **359**, 1170–1177

69. Mateus, A., Kurzawa, N., Becher, I., Sridharan, S., Helm, D., Stein, F., Typas, A., and Savitski, M. M. (2020) Thermal proteome profiling for interrogating protein interactions. Mol. Syst. Biol. 10.15252/MSB.20199232

70. Hashimoto, Y., Sheng, X., Murray-Nerger, L. A., and Cristea, I. M. (2020) Temporal dynamics of protein complex formation and dissociation during human cytomegalovirus infection. Nat. Commun. 10.1038/s41467-020-14586-5

71. Winkler, I., Marx, A. D., Lariviere, D., Heinze, R. J., Cristovao, M., Reumer, A., Curth, U., Sixma, T. K., and Friedhoff, P. (2011) Chemical trapping of the dynamic MutS-MutL complex formed in DNA mismatch repair in Escherichia coli. Journal of Biological Chemistry. 286, 17326–17337

72. Groothuizen, F. S., Winkler, I., Cristóvão, M., Fish, A., Winterwerp, H. H., Reumer, A., Marx, A. D., Hermans, N., Nicholls, R. A., Murshudov, G. N., Lebbink, J. H., Friedhoff, P., and Sixma, T. K. (2015) MutS/MutL crystal structure reveals that the MutS sliding clamp loads MutL onto DNA. Elife. 10.7554/ELIFE.06744

73. Lee, J. M., Ramos, E. M., Lee, J. H., Gillis, T., Mysore, J. S., Hayden, M. R., Warby, S. C., Morrison, P., Nance, M., Ross, C. A., Margolis, R. L., Squitieri, F., Orobello, S., Di Donato, S., Gomez-Tortosa, E., Ayuso, C., Suchowersky, O., Trent, R. J. A., McCusker, E., Novelletto, A., Frontali, M., Jones, R., Ashizawa, T., Frank, S., Saint-Hilaire, M. H., Hersch, S. M., Rosas, H. D., Lucente, D., Harrison, M. B., Zanko, A., Abramson, R. K., Marder, K., Sequeiros, J., Paulsen, J. S., Landwehrmeyer, G. B., Myers, R. H., MacDonald, M. E., and Gusella, J. F. (2012) CAG repeat expansion in Huntington disease determines age at onset in a fully dominant fashion. Neurology. 78, 690–695

74. Flower, M., Lomeikaite, V., Ciosi, M., Cumming, S., Morales, F., Lo, K., Hensman Moss, D., Jones, L., Holmans, P., Monckton, D. G., Tabrizi, S. J., Kraus, P., Hoffman, R., Tobin, A., Borowsky, B., Keenan, S., Whitlock, K. B., Queller, S., Campbell, C., Wang, C., Langbehn, D., Axelson, E., Johnson, H., Acharya, T., Cash, D. M., Frost, C., Jones, R., Jurgens, C., Hart, E. P. T., Van Der Grond, J., Witjes-Ane, M. N. N., Roos, R. A. C., Dumas, E. M., Van Den Bogaard, S. J. A., Stopford, C., Craufurd, D., Callaghan, J., Arran, N., Rosas, D. D., Lee, S., Monaco, W., O’Regan, A., Milchman, C., Frajman, E., Labuschagne, I., Stout, J., Campbell, M., Andrews, S. C., Bechtel, N., Reilmann, R., Bohlen, S., Kennard, C., Berna, C., Hicks, S., Durr, A., Pourchot, C., Bardinet, E., Nigaud, K., Valabrègue, R., Lehericy, S., Marelli, C., Jauffret, C., Justo, D., Leavitt, B., Decolongon, J., Sturrock, A., Coleman, A., Dar Santos, R., Patel, A., Gibbard, C., Whitehead, D., Wild, E., Owen, G., Crawford, H., Malone, I., Lahiri, N., Fox, N. C., Hobbs, N. Z., Scahill, R. I., Ordidge, R., Pepple, T., Read, J., Say, M. J., Landwehrmeyer, B., Daidj, F., Bassez, G., Lignier, B., Couppey, F., Delmas, S., Deux, J. F., Hankiewicz, K., Dogan, C., Minier, L., Chevalier, P., Hamadouche, A., Catt, M., Van Hees, V., Catt, S., Schwalber, A., Dittrich, J., Kierkegaard, M., Wenninger, S., Schoser, B., Schüller, A., Stahl, K., Künzel, H., Wolff, M., Jellinek, A., Moreno, C. J., Gorman, G., Lochmüller, H., Trenell, M., Van Laar, S., Wood, L., Cassidy, S., Newman, J., Charman, S., Steffaneti, R., Taylor, L., Brownrigg, A., Day, S., Atalaia, A., Raaphorst, J., Okkersen, K., Van Engelen, B., Nikolaus, S., Cornelissen, Y., Van Nimwegen, M., Maas, D., Klerks, E., Bouman, S., Knoop, H., Heskamp, L., Heerschap, A., Rahmadi, R., Groot, P., Heskes, T., Kapusta, K., Glennon, J., Abghari, S., Aschrafi, A., Poelmans, G., Treweek, S., Hogarth, F., Littleford, R., Donnan, P., Hapca, A., Hannah, M., Mckenzie, E., Rauchhaus, P., Cumming, S. A., Adam, B., Faber, C., and Merkies, I. (2019) MSH3 modifies somatic instability and disease severity in Huntington’s and myotonic dystrophy type 1. Brain. 142, 1876

75. Couch, Y. (2023) Challenges associated with using extracellular vesicles as biomarkers in neurodegenerative disease. Expert Rev. Mol. Diagn. 23, 1091–1105

76. Brás, I. C., Xie, Y., and Southwell, A. L. (2025) Neuron-derived extracellular vesicles in plasma present a potential non-invasive biomarker for Huntingtin protein and RNA assessment in Huntington disease. bioRxiv. 10.1101/2025.07.17.665403

77. Belgrad, J., Greco, T. M., Sapp, E., Summers, A., O’Reilly, D., Luu, E., Hutton, J. E., Yamada, N., Fakih, H. H., Furgal, R., Echeverria, D., McHugh, N., Bramato, B., Furguson, C., Hildebrand, S., Allen, S., Gaston, N., Cooper, D., Maebius, A., Gross, K. Y., Vogt, T. F., Finley, M., Prasad, B., DiFiglia, M., Cristea, I. M., Aronin, N., and Khvorova, A. (2026) Mismatch repair dissection by in vivo RNAi reveals dose-dependent modulators of somatic instability and proteome remodeling in Huntington’s disease. bioRxiv. 10.64898/2026.06.19.733435

78. An, S., and Fu, L. (2018) Small-molecule PROTACs: An emerging and promising approach for the development of targeted therapy drugs. EBioMedicine. 36, 553–562

79. Tomé, S., Simard, J. P., Slean, M. M., Holt, I., Morris, G. E., Wojciechowicz, K., te Riele, H., and Pearson, C. E. (2013) Tissue-specific mismatch repair protein expression: MSH3 is higher than MSH6 in multiple mouse tissues. DNA Repair (Amst). 12, 46–52

80. Carroll, K. M., Simpson, D. M., Eyers, C. E., Knight, C. G., Brownridge, P., Dunn, W. B., Winder, C. L., Lanthaler, K., Pir, P., Malys, N., Kell, D. B., Oliver, S. G., Gaskell, S. J., and Beynon, R. J. (2011) Absolute Quantification of the Glycolytic Pathway in Yeast: *Molecular & Cellular Proteomics*. 10, M111.007633

81. Takemori, N., Takemori, A., Tanaka, Y., Endo, Y., Hurst, J. L., Gomez-Baena, G., Harman, V. M., and Beynon, R. J. (2017) MEERCAT: Multiplexed Efficient Cell Free expression of recombinant qconcats for large scale absolute proteome quantification. Molecular and Cellular Proteomics. 16, 2169–2183

82. Narumi, R., Shimizu, Y., Ukai-Tadenuma, M., Ode, K. L., Kanda, G. N., Shinohara, Y., Sato, A., Matsumoto, K., and Ueda, H. R. (2016) Mass spectrometry-based absolute quantification reveals rhythmic variation of mouse circadian clock proteins. Proc. Natl. Acad. Sci. U. S. A. 113, E3461–E3467

83. Brownridge, P., Holman, S. W., Gaskell, S. J., Grant, C. M., Harman, V. M., Hubbard, S. J., Lanthaler, K., Lawless, C., O’cualain, R., Sims, P., Watkins, R., and Beynon, R. J. (2011) Global absolute quantification of a proteome: Challenges in the deployment of a QconCAT strategy. Proteomics. 11, 2957–2970

84. Benesova, E., Vidova, V., and Spacil, Z. (2021) A comparative study of synthetic winged peptides for absolute protein quantification. Scientific Reports |. 11, 10880

85. Xian, F., Zi, J., Wang, Q., Lou, X., Sun, H., Lin, L., Hou, G., Rao, W., Yin, C., Wu, L., Li, S., and Liu, S. (2016) Peptide biosynthesis with stable isotope labeling from a cell-free expression system for targeted proteomics with absolute quantification. Molecular and Cellular Proteomics. 15, 2819–2828

86. Han, S. H., Kim, J. S., Lee, Y., Choi, H., Kim, J. W., Na, D. L., Yang, E. G., Yu, M. H., Hwang, D., Lee, C., and Mook-Jung, I. (2014) Both targeted mass spectrometry and flow sorting analysis methods detected the decreased serum apolipoprotein e level in alzheimer’s disease patients. Molecular and Cellular Proteomics. 13, 407–419

87. Pannee, J., Blennow, K., Zetterberg, H., and Portelius, E. (2017) Video article absolute quantification of Aβ1-42 in CSF using a mass spectrometric reference measurement procedure. Journal of Visualized Experiments. 10.3791/55386

88. Rzagalinski, I., Rao, V., Bogdanova, A., Hersemann, L., and Shevchenko, A. (2025) Targeted Absolute Quantification of Protein Biomarkers in Cerebrospinal Fluid by FastCAT. Methods in Molecular Biology. 2914, 65–74

89. Bader, J. M., Albrecht, V., and Mann, M. (2023) MS-Based Proteomics of Body Fluids: The End of the Beginning. Molecular and Cellular Proteomics. 10.1016/j.mcpro.2023.100577

90. Zahn, E., Xie, Y., Liu, X., Karki, R., Searfoss, R. M., de Luna Vitorino, F. N., Lempiäinen, J. K., Gongora, J., Lin, Z., Zhao, C., Yuan, Z. F., and Garcia, B. A. (2025) Development of a High-Throughput Platform for Quantitation of Histone Modifications on a New QTOF Instrument. Molecular and Cellular Proteomics. 10.1016/j.mcpro.2024.100897

91. Garofalo, R., Wohlgemuth, I., Pearson, M., Lenz, C., Urlaub, H., and Rodnina, M. V. (2019) Broad range of missense error frequencies in cellular proteins. Nucleic Acids Res. 47, 2932–2945

92. Tang, M. C. W., Binos, S., Ong, E. K., Wong, L. H., and Mann, J. R. (2014) High histone variant H3.3 content in mouse prospermatogonia suggests a role in epigenetic reformatting. Chromosoma. 123, 587–595

93. Smits, A. H., Jansen, P. W. T. C., Poser, I., Hyman, A. A., and Vermeulen, M. (2013) Stoichiometry of chromatin-associated protein complexes revealed by label-free quantitative mass spectrometry-based proteomics. Nucleic Acids Res. 10.1093/nar/gks941

94. Wohlgemuth, I., Lenz, C., and Urlaub, H. (2015) Studying macromolecular complex stoichiometries by peptide-based mass spectrometry. Proteomics. 15, 862–879

95. Wheeler, V. C., Gutekunst, C. A., Vrbanac, V., Lebel, L. A., Schilling, G., Hersch, S., Friedlander, R. M., Gusella, J. F., Vonsattel, J. P., Borchelt, D. R., and MacDonald, M. E. (2002) Early phenotypes that presage late-onset neurodegenerative disease allow testing of modifiers in Hdh CAG knock-in mice. Hum. Mol. Genet. 11, 633–640

96. Wheeler, V. C., White, J. K., Gutekunst, C. A., Vrbanac, V., Weaver, M., Li, X. J., Li, S. H., Yi, H., Vonsattel, J. P., Gusella, J. F., Hersch, S., Auerbach, W., Joyner, A. L., and MacDonald, M. E. (2000) Long glutamine tracts cause nuclear localization of a novel form of huntingtin in medium spiny striatal neurons in Hdh(Q92) and Hdh(Q111) knock-in mice. Hum. Mol. Genet. 9, 503–513

97. Carty, N., Berson, N., Tillack, K., Thiede, C., Scholz, D., Kottig, K., Sedaghat, Y., Gabrysiak, C., Yohrling, G., Von Der Kammer, H., Ebneth, A., Mack, V., Munoz-Sanjuan, I., and Kwak, S. (2015) Characterization of HTT inclusion size, location, and timing in the zQ175 mouse model of Huntington’s disease: An in vivo high-content imaging study. PLoS One. 10.1371/JOURNAL.PONE.0123527,

98. Smith, E. J., Sathasivam, K., Landles, C., Osborne, G. F., Mason, M. A., Gomez-Paredes, C., Taxy, B. A., Milton, R. E., Ast, A., Schindler, F., Zhang, C., Duan, W., Wanker, E. E., and Bates, G. P. (2023) Early detection of exon 1 huntingtin aggregation in zQ175 brains by molecular and histological approaches. Brain Commun. 10.1093/BRAINCOMMS/FCAD010,

99. Desmond, Atwal, R. S., Xia, J., and Truant, R. (2012) Identification of a karyopherin β1/β2 proline-tyrosine nuclear localization signal in huntingtin protein. Journal of Biological Chemistry. 287, 39626–39633

100. De Rooij, K. E., Dorsman, J. C., Smoor, M. A., Den Dunnen, J. T., and Van Ommen, G. J. B. (1996) Subcellular localization of the Huntington’s disease gene product in cell lines by immunofluorescence and biochemical subcellular fractionation. Hum. Mol. Genet. 5, 1093–1099

101. Atwal, R. S., Xia, J., Pinchev, D., Taylor, J., Epand, R. M., and Truant, R. (2007) Huntingtin has a membrane association signal that can modulate huntingtin aggregation, nuclear entry and toxicity. Hum. Mol. Genet. 16, 2600–2615

102. Panov, A. V., Gutekunst, C. A., Leavitt, B. R., Hayden, M. R., Burke, J. R., Strittmatter, W. J., and Greenamyre, J. T. (2002) Early mitochondrial calcium defects in Huntington’s disease are a direct effect of polyglutamines. Nat. Neurosci. 5, 731–736

103. Grima, J. C., Daigle, J. G., Arbez, N., Cunningham, K. C., Zhang, K., Ochaba, J., Geater, C., Morozko, E., Stocksdale, J., Glatzer, J. C., Pham, J. T., Ahmed, I., Peng, Q., Wadhwa, H., Pletnikova, O., Troncoso, J. C., Duan, W., Snyder, S. H., Ranum, L. P. W., Thompson, L. M., Lloyd, T. E., Ross, C. A., and Rothstein, J. D. (2017) Mutant Huntingtin Disrupts the Nuclear Pore Complex. Neuron. 94, 93–107.e6

104. Truant, R., Atwal, R. S., and Burtnik, A. (2007) Nucleocytoplasmic trafficking and transcription effects of huntingtin in Huntington’s disease. Prog. Neurobiol. 83, 211–227

105. Duyao, M. P., Auerbach, A. B., Ryan, A., Persichetti, F., Barnes, G. T., McNeil, S. M., Ge, P., Vonsattel, J. P., Gusella, J. F., Joyner, A. L., and MacDonald, M. E. (1995) Inactivation of the mouse Huntington’s disease gene homolog Hdh. Science. 269, 407–410

106. Zeitlin, S., Liu, J. P., Chapman, D. L., Papaioannou, V. E., and Efstratiadis, A. (1995) Increased apoptosis and early embryonic lethality in mice nullizygous for the Huntington’s disease gene homologue. Nat. Genet. 11, 155–163

107. Steffan, J. S., Kazantsev, A., Spasic-Boskovic, O., Greenwald, M., Zhu, Y. Z., Gohler, H., Wanker, E. E., Bates, G. P., Housman, D. E., and Thompson, L. M. (2000) The Huntington’s disease protein interacts with p53 and CREB-binding protein and represses transcription. Proc. Natl. Acad. Sci. U. S. A. 97, 6763–6768

108. Zuccato, C., Tartari, M., Crotti, A., Goffredo, D., Valenza, M., Conti, L., Cataudella, T., Leavitt, B. R., Hayden, M. R., Timmusk, T., Rigamonti, D., and Cattaneo, E. (2003) Huntingtin interacts with REST/NRSF to modulate the transcription of NRSE-controlled neuronal genes. Nat. Genet. 35, 76–83

109. Seong, I. S., Woda, J. M., Song, J. J., Lloret, A., Abeyrathne, P. D., Woo, C. J., Gregory, G., Lee, J. M., Wheeler, V. C., Walz, T., Kingston, R. E., Gusella, J. F., Conlon, R. A., and MacDonald, M. E. (2010) Huntingtin facilitates polycomb repressive complex 2. Hum. Mol. Genet. 19, 573–583

110. Kozłowska, E., Ciołak, A., Adamek, G., Szcześniak, J., and Fiszer, A. (2025) HTT loss-of-function contributes to RNA deregulation in developing Huntington’s disease neurons. Cell Biosci. 10.1186/S13578-025-01443-5

111. Sun, X., Liu, L., Wu, C., Li, X., Guo, J., Zhang, J., Guan, J., Wang, N., Gu, L., Yang, X. W., and Li, G. M. (2024) Mutant huntingtin protein induces MLH1 degradation, DNA hyperexcision, and cGAS-STING-dependent apoptosis. Proc. Natl. Acad. Sci. U. S. A. 10.1073/PNAS.2313652121

112. Fishwick, K. M., Gomez Vieito, D., Greco, G., Collotta, G., Gatti, M., Kulik, A. A., Guérois, R., Corbeski, I., Phadte, A. S., Senoussi, I., Cejka, P., Pluciennik, A., Porro, A., and Sartori, A. A. (2025) Disruption of protein-protein interaction hotspots in the C-terminal domain of MLH1 confers mismatch repair deficiency. NAR Cancer. 7, zcaf055

113. Redon, S., Zemp, I., and Lingner, J. (2013) A three-state model for the regulation of telomerase by TERRA and hnRNPA1. Nucleic Acids Res. 41, 9117–9128

114. Simon, M. N., Dubrana, K., and Palancade, B. (2024) On the edge: how nuclear pore complexes rule genome stability. Curr. Opin. Genet. Dev. 84, 102150

115. Rousseau, V., Einig, E., Jin, C., Horn, J., Riebold, M., Poth, T., Jarboui, M. A., Flentje, M., and Popov, N. (2023) Trim33 masks a non-transcriptional function of E2f4 in replication fork progression. Nat. Commun. 14, 5143

116. Haye, J. E., and Gammie, A. E. (2015) The Eukaryotic Mismatch Recognition Complexes Track with the Replisome during DNA Synthesis. PLoS Genet. 11, e1005719

